# Low diversity and instability of the sinus microbiota over time in adults with cystic fibrosis

**DOI:** 10.1101/2022.01.18.476814

**Authors:** Catherine R. Armbruster, Kelvin Li, Megan R. Kiedrowski, Anna C. Zemke, Jeffrey A. Melvin, John Moore, Samar Atteih, Adam C. Fitch, Matthew DuPont, Christopher D. Manko, Madison L. Weaver, Jordon R. Gaston, John F. Alcorn, Alison Morris, Barbara A. Methé, Stella E. Lee, Jennifer M. Bomberger

## Abstract

**Background:** Chronic rhinosinusitis (CRS) is a common, yet underreported and understudied manifestation of upper respiratory disease in people with cystic fibrosis (CF). There are currently no standard of care guidelines for the management of CF CRS, but treatment of upper airway disease may ameliorate lower airway disease. We sought to inform future treatment guidelines by determining whether changes to sinus microbial community diversity and specific taxa known to cause CF lung disease are associated with increased respiratory disease and inflammation.

**Methods:** We performed 16S rRNA gene sequencing, supplemented with cytokine analyses, microscopy, and bacterial culturing, on samples from the sinuses of 27 adults with CF CRS at the University of Pittsburgh’s CF Sinus Clinic. At each study visit, participants underwent endoscopic paranasal sinus sampling and clinical evaluation. We identified key drivers of microbial community composition and evaluated relationships between diversity and taxa with disease outcomes and inflammation.

**Findings:** Sinus community diversity was low and the composition was unstable, with many participants exhibiting alternating dominance between *Pseudomonas aeruginosa* and Staphylococci over time. Despite a tendency for dominance by these two taxa, communities were highly individualized and shifted composition during exacerbation of sinus disease symptoms. Exacerbations were also associated with communities dominated by *Staphylococcus* spp. Reduced microbial community diversity was linked to worse sinus disease and the inflammatory status of the sinuses (including increased IL-1*β*). Increased IL-1*β* was also linked to worse sinus endoscopic appearance, and other cytokines were linked to microbial community dynamics.

**Interpretation:** To our knowledge, this is the largest longitudinal study of microbial communities and cytokine secretion in CF CRS. Our work revealed previously unknown instability of sinus microbial communities and a link between inflammation, lack of microbial community diversity, and worse sinus disease.

**Funding:** Cystic Fibrosis Foundation and US National Institutes of Health.

**Research in Context:** *Evidence before this study:* A search of the PubMed database on October 11, 2021 with the terms [cystic fibrosis sinus microbiome] yielded 16 results, and we have identified seven primary research articles on the CF CRS microbiome (including re-analyses of existing datasets). Most are cross-sectional cohort analyses, along with one prior longitudinal study of four adults at the University of Auckland, New Zealand. Together, these prior studies reveal similarities between CF CRS and CF sputum microbiomes, including low community diversity associated with sinus disease, the presence of common CF-associated microbes in the sinuses, and prevalence of sinus communities dominated by *P. aeruginosa* or *Staphylococcus aureus*. High levels of IL-1*β* are linked to the presence of nasal polyps in CF CRS, and polymorphisms in the IL-1 receptor antagonist gene are associated with risk of CRS outside of the context of CF. Two prior studies of this cohort have been performed by our laboratory. One describes clinical indicators of CF sinus disease and the other links sinus infection biogeography to *P. aeruginosa* evolutionary genomics.

*Added value of this study:* Our study is the first to examine longitudinal relationships between the host immune response (through cytokine profiling) and microbiota dynamics in CF CRS, including linking elevated IL-1*β* to worse sinus disease through reduced sinus microbial community diversity. The longitudinal nature of our study also allowed us to uncover striking temporal instability of microbial communities in approximately half of our cohort’s sinus microbial communities over two years, including switching between communities dominated by *P. aeruginosa* and *Staphylococcus* spp. This instability could hinder attempts to link the relative abundance of taxa to clinical outcomes of interest in cross-sectional studies (e.g., markers of disease progression). We also identified patterns of synergy and antagonism between specific taxa, and impacts of the host immune response in the sinuses on community composition.

*Implications of all the available evidence:* Together with prior CF CRS microbiome studies, our study underscores similarities between sinus and lower respiratory tract microbial community structure in CF, and we show how community structure tracks with inflammation and several disease measures. This work strongly suggests that clinical management of CRS could be leveraged to improve overall respiratory health in CF. Our work implicates elevated IL-1*β* in reduced microbiota diversity and worse sinus disease in CF CRS, suggesting applications for existing therapies targeting IL-1*β*. Finally, the widespread use of highly effective CFTR modulator therapy has led to less frequent availability of spontaneous expectorated sputum for microbiological surveillance of lung infections. A better understanding of CF sinus microbiology could provide a much-needed alternative site for monitoring respiratory infection status by important CF pathogens.

## Introduction

The upper airways are constantly exposed to microbes inhaled from the environment. In healthy individuals, these microbes are captured by mucus produced by the sinonasal epithelium and removed by mucocilliary clearance, but this process is impaired in people with the genetic disorder cystic fibrosis (CF).^1^ The sinonasal cavity is thought to be the first site in the respiratory tract to be colonized by opportunistically pathogenic microbes that may seed downstream lung disease in CF.^2^ Chronic rhinosinusitis (CRS), defined as symptomatic chronic infection and inflammation of the sinonasal cavity, is common among people with CF, yet under-reported and the interactions between microbes in the upper respiratory tract, local inflammatory responses, and clinical outcomes are poorly understood. The unified airway hypothesis is a conceptual framework originating in the field of asthma research that links upper (URT) and lower (LRT) respiratory tract disease.^3^ This framework proposes that treatment of URT symptoms can improve LRT disease and vice-a-versa. A growing body of literature supports similarities and interplay between CRS and LRT disease in CF. For example, the microbiota of CF CRS resembles that of the LRT in terms of taxa present and diversity^4–6;^ children harbor comparatively diverse microbes, whereas adults tend to be dominated by one or very few organisms.^7^ Furthermore, medical or surgical management of sinus disease symptoms may lead to better LRT outcomes in CF.^8, 9^ Recently, studies by our team and others have shown that evolved traits and evolutionary strategies of *Pseudomonas aeruginosa* isolated from the sinuses of people with CF CRS resemble those previously reported among CF lung populations.^10, 11^ We have also shown that sinus exacerbation increases the odds of a subsequent pulmonary exacerbation, and recently others have shown that sinonasal quality of life worsens during CF LRT exacerbations.^12, 13^ Together, these studies strongly suggest that CF CRS impacts LRT disease, yet there are currently no standard of care guidelines for the management of CF CRS. One gap in the CF CRS literature is our lack of understanding regarding whether and how CF sinus communities change over time and as the conditions in the surrounding host environment change (e.g., during periods of increased inflammation and/or exacerbation of symptoms).

The goal of this study was to evaluate how the microbial composition and inflammatory environment of the sinuses relates to upper and lower airway disease in adults with CF CRS. We hypothesized that lack of sinus microbial community diversity and changes in relative abundance of opportunistic pathogens or pathobionts (commensals that can cause disease under certain circumstances) would be associated with increased sinus disease severity and inflammation. In addition to revealing similarities between microbiota-related correlates of CF sinus disease and inflammation to those described for the lower respiratory tract in CF, our study hints at potential new therapeutic opportunities based on these microbe-immune interactions and highlights the relevance of sinus disease to overall CF respiratory health.

## Results

### Cohort demographics and association of CF-related diabetes (CFRD) with lower respiratory disease

We performed a longitudinal study of 33 adults with CF and symptomatic CRS who had undergone prior functional endoscopic sinus surgery (FESS) as treatment for CF CRS (Table 1).^13^ During quarterly clinic visits and unscheduled visits due to exacerbation of clinical symptoms, we obtained at least one endoscopically guided specimen for 16S amplicon sequencing from 27 of the 33 study participants (longitudinal microbiota samples collected from 18 of the 27). Additional samples included sinus secretions that were collected for inflammatory cytokine analyses, bacterial culturing, and microscopy. The following demographic and clinical characteristics of the cohort (covariates) were controlled for in most of the later analyses: Patient ID (to control for repeated measures within study participants), age, sex, CFTR mutation class, diagnosis of CF-related diabetes (CFRD), BMI, and topical antibiotic use. We tested for associations between these covariates and patient outcomes examined throughout this study (sinus exacerbation, pulmonary exacerbation, sinus disease score [Sino-nasal Outcome Test, SNOT-22^14^; modified Lund-Kennedy, mLK^15^], and lung function [FEV1]), independent of information on the microbiota. We found that CFRD was associated with reduced FEV1 (univariate regression coefficient: -68.3, Holm-Bonferroni adjusted p < 0.05). No other covariates were significantly associated with the outcomes of interest.

**Table 1.** Demographics of the adult CF CRS microbiota study cohort. The cohort includes 27 people from the larger 33 person study, for whom we sequenced at least one 16S amplicon microbiota sample from a paranasal sinus swab collected by endoscope. For 18 of these 27 people, we sequenced at least two samples, giving us longitudinal information. Drug use is reported for any time during study. CF-related diabetes (CFRD) is reported for at any time within the study or +/- 12 months of enrollment. Clinical parameters of the full cohort (N = 33) were published in Zemke, IFAR 2019.

|  | Microbiota cohort (N = 27/33) |
| --- | --- |
| Longitudinal microbiota samples (%) | 18/27 (66.6) |
| Median age on enrollment (range) | 27.6 (19.7-43.6) |
| Male (%) | 9/27 (33.3) |
| CFTR genotype (%) | ΔF508 homozygous 13/27 (48.2) |
|  | ΔF508/other 11/27 (40.7) |
|  | Other/other 2/27 (7.4) |
|  | Missing 1/27 (3.7) |
| CFRD (%) | 12/27 (44.4) |
| Topical sinus antibiotic use? (%) | 22/27 (81.5) |
| Topical or oral steroid use? (%) | 21/27 (77.7) |
| CFTR corrector/modulator use? (%) | 10/27 (37) |

### Low diversity and instability of sinus microbial communities in adults with CF CRS

Sinus microbial community diversity in our cohort was low, with the median Shannon diversity from all participants being 0.35 (IQR 0.11-0.62) and Simpson 0.18 (IQR 0.04-0.86; Figure 1A). The median evenness was 0.15 (IQR 0.04-0.63), suggesting communities were dominated by a subset of the taxa present. Finally, the Tail statistic (“τ”) is a rank-based diversity measure that is more sensitive to changes in low-abundance taxa. The median τ was 0.46 (IQR 0.45-0.66) , but had a fairly large range (from 0.04-5.6), indicating that diverse low-abundance taxa are present in the sinuses of some, but not all, study participants (Supplemental table 1). We detected a total of 302 genera (Supplemental table 2).

**Figure 1.**
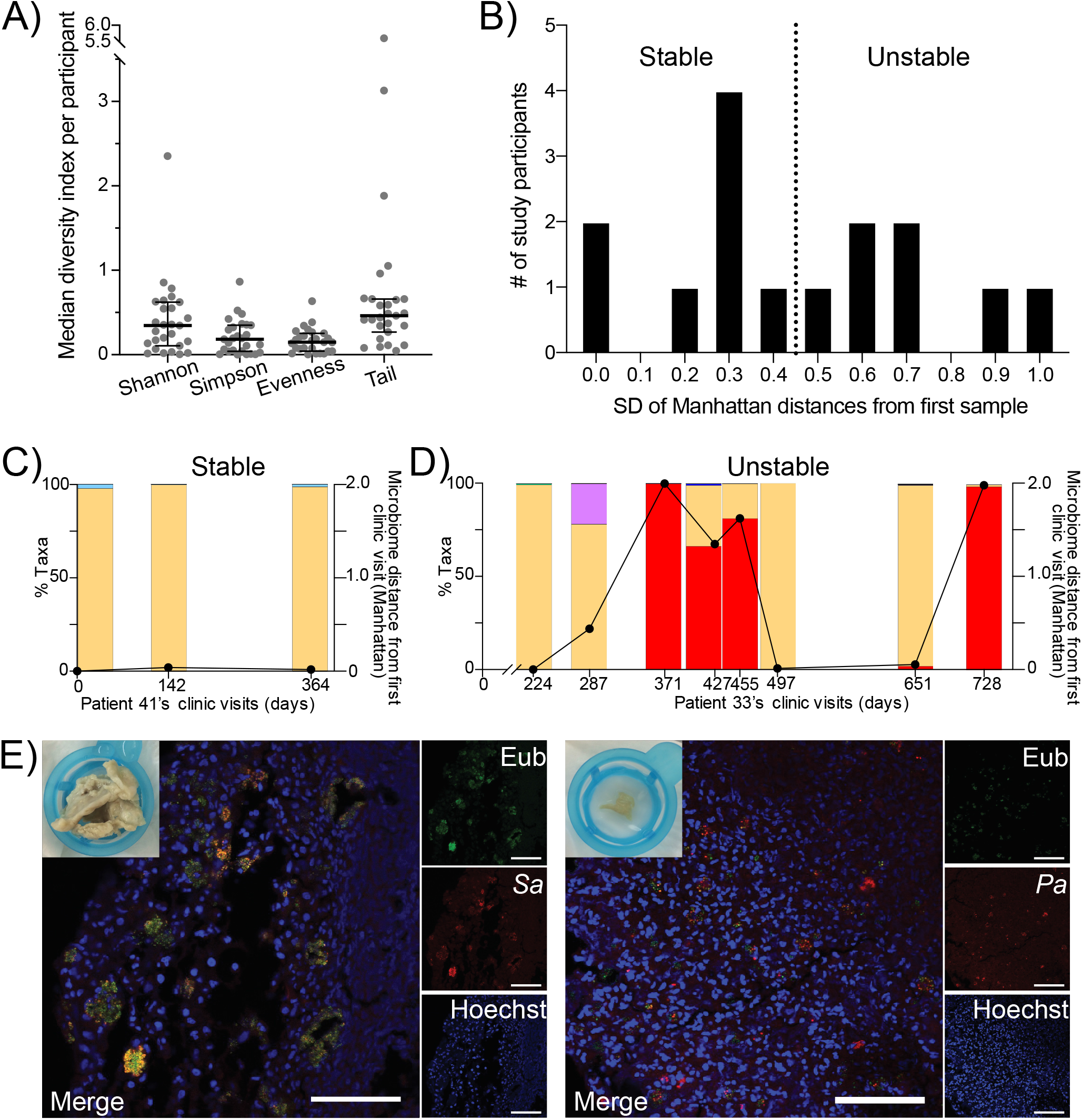
Sinus microbial community diversity is low and the composition can be unstable in adults with CF chronic rhinosinusitis, with most study participants’ sinuses dominated by *P. aeruginosa* or *Staphylococcus* spp. A) Boxplots depicting the median Shannon or Simpson diversity indices, evenness, or Tail statistic per study participant. Overlaid are the cohort’s median and interquartile range. N = 27 participants. B) Study participants’ sinus microbial communities were categorized as being relative stable or unstable over time. Histogram binning of study participants based on the standard deviation (SD) of each of their study visits’ Manhattan distances from the first microbiota sample sequenced. The dotted line indicates the median SD across all participants. N = 15 study participants with 3 or more visits. C) An example of a study participant (Patient 41), categorized as relatively stable based on the results in Figure 1B, whose sinus microbiota was consistently dominated by *P. aeruginosa* over time. Taxonomic bar plots depict the relative abundance of *Pseudomonadaceae* (*P. aeruginosa*; peach). The overlaid line plotted on the right y-axis is the Manhattan distance of each sample from their first microbiota sample. D) An example of a study participant (Patient 33), categorized as unstable based on the results in Figure 1B, who exhibited switching between *P. aeruginosa* and Staphylococcus spp. dominance over time. Taxa bar plots depicting the relative abundance of *Pseudomonadaceae* (*P. aeruginosa*; peach), *Staphylococcus* spp. (red), *Corynebacterium* spp. (purple), and other low abundance taxa. The overlaid line plotted on the right y axis is the Manhattan distance of each sample from their first microbiota sample. E) Representative FISH images of CF CRS microbial communities from explanted obstructive sinus debris sampled by endoscope from two study participants. On the left, red = *S. aureus* probe, green = Eubacterial (universal) probe, blue = Hoechst stain (mostly host cell nuclei). On the right, red = *P. aeruginosa*, green = Eubacterial (universal) probe, blue = Hoechst stain. Scale bars = 50 μm. The macroscopic image at the top left of each FISH image depicts the mucopurulent sinus sample prior to processing for microscopy.

While diversity indices of the sequenced sinus microbiotas were low, microbial community composition was unstable for many study participants (Figure 1B). For individuals that contributed at least 3 longitudinal microbiota samples, we quantified this instability based on the standard deviation of the microbiota distance (Manhattan) at later timepoints relative to the first timepoint. The histogram of these values was bimodal on either side of the median value for the study cohort, allowing us to classify eight individuals as relatively stable (for example, the individual whose taxonomic barplots are shown in Figure 1C) and seven as relatively unstable (for example, in Figure 1D). The most common bacterial taxa were *Staphylococcus* spp. and *Pseudomonadaceae* (Supplemental table 2), and the relative abundances of these two taxa varied over time in several study participants from whom we collected longitudinal samples (Figure 1D and Supplemental figure 1). Based on Sanger sequencing of the 16S rRNA gene following bacterial culture, the *Pseudomonadaceae* taxon represents *Pseudomonas aeruginosa* and is referred to as such hereafter. We then examined the biogeography of *P. aeruginosa* and/or *S. aureus* in these communities, using fluorescent *in* situ hybridization (FISH) on explanted obstructive sinus material that was surgically debrided as part of routine clinical care. We found that both *P. aeruginosa* and *S. aureus* reside as small aggregated communities in close association with host cells in the sinuses (Figure 1E). Eubacterial labeling did not fully overlap with species-specific probes, suggesting other unidentified species were present in mixed-species aggregates. Overall, these results suggest that while CF CRS microbes can reside in small, sparse aggregates where diversity is low, the aggregates can contain mixed species in proximity with each other and with host cells. Furthermore, the overall taxonomic composition of sinus communities can be rather unstable, especially in individuals co-infected by *P. aeruginosa* and *Staphylococcus* spp.

While *P. aeruginosa* and *Staphylococcus* spp. were abundant in many study participants, other microbes were stably present as well. Regarding patterns of co-occurrence among microbes, we identified positive correlations between the presence of *Corynebacterium* spp. and *Dolosigranulum* spp., whereas *P. aeruginosa* and *Burkholderia* spp. exhibited an antagonistic relationship (Supplemental figure 2). In addition to taxa recognized as members of the nasal, sinus, or oral microbiotas of healthy adults, we identified bacteria known to be present in potable water and capable of causing opportunistic infections in susceptible populations (e.g. *Sphingomonas* spp. in 11 out of 27 participants, *Bradyrhizobium* spp. in ten, *Methylobacterium* spp. in nine, and *Delftia* spp. in six; Supplemental table 2).^16^ The composition of environmental and reagent control samples processed and sequenced alongside our study specimens was distinct from clinical specimens (PERMANOVA; p < 0.0001), and the controls had significantly lower read counts compared to the study samples (T-test; p-value <0.0001). These controls suggest that the drinking water taxa were not due to contamination of clinical specimens. Overall, these findings demonstrate that while most sinus communities were dominated by *P. aeruginosa* and/or *Staphylococcus* spp., a variety of other taxa were also detected. The presence of bacteria frequently reported to be present in potable water suggests a potential exposure route of the sinuses to opportunistically pathogenic microbes that contribute to diversity of low-abundance taxa.

**Figure 2.**
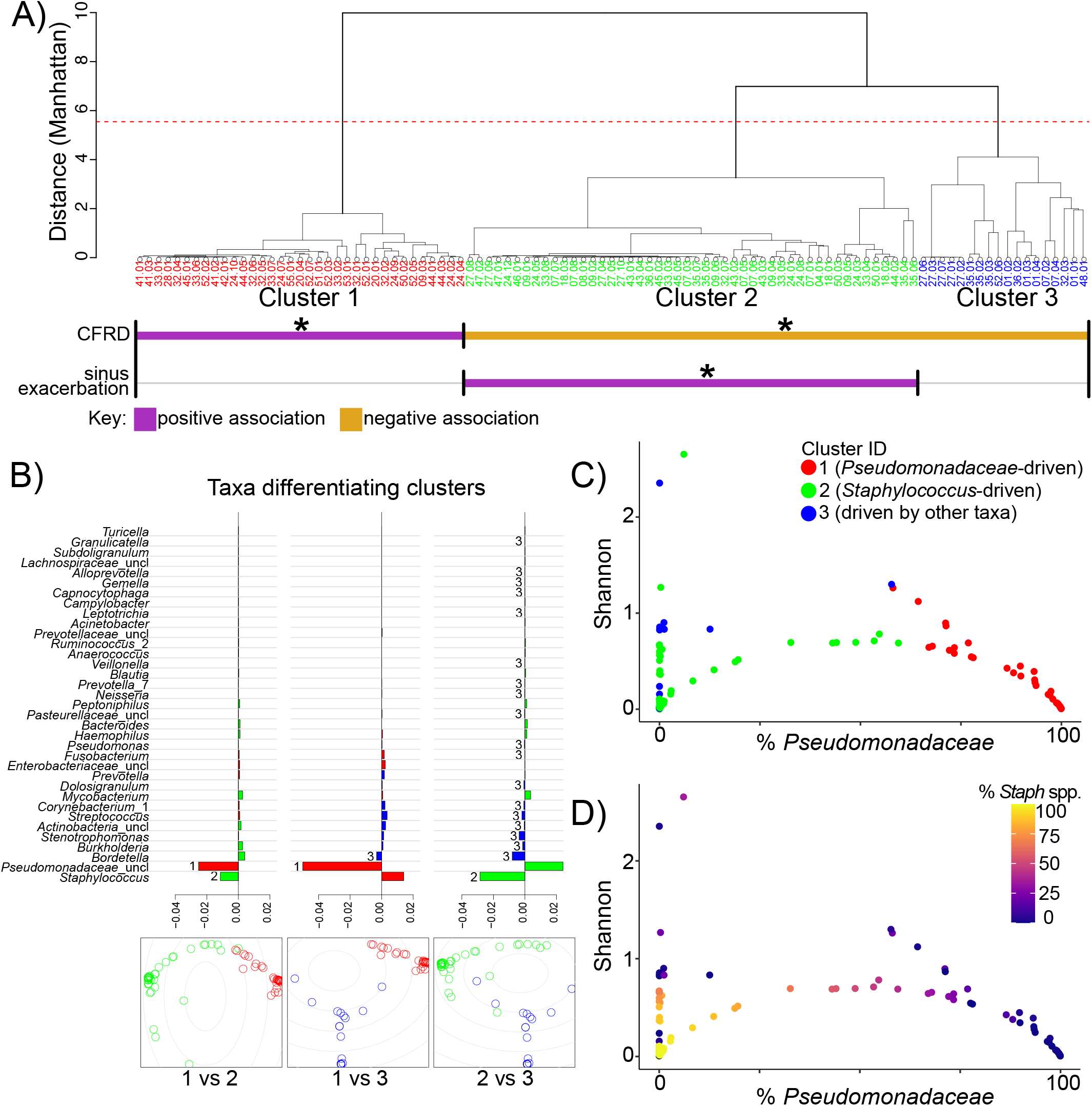
Relative abundance of *Staphylococcus* spp. or *P. aeruginosa* almost exclusively drives low microbial diversity and community structure. A) Dendrogram depicting individual patient samples hierarchically clustered using Ward’s minimum variance method on the inter- sample Manhattan distance. Individual microbiota samples are colored based on their cluster (1 = red, 2 = green, 3 = blue). Covariates or sinus disease outcome measures that significantly correlated with cluster membership by multinomial linear regression are summarized below the clusters. CFRD is positively associated with Cluster 1 and negatively associated with Clusters 2 and 3. Sinus exacerbation is positively associated with Cluster 2. B) Identifying which and to what extent each taxon drives the differences between clusters was calculated by comparing the coefficient of determination (R^2^) for a reduced model (without a taxon of interest) against a full model (with all taxa included) with the R^2^ ratio: R^2^ reduced / R^2^ full. If excluding a taxon (evaluated with the reduced model) reduces the separation between two clusters relative to keeping it (evaluated with the full model), then it was an important taxon to the cluster, and the R^2^ ratio would be <1. Values plotted are log10(R^2^ reduced / R^2^ full), with negative values indicating the most influential taxa separating the two clusters compared. The bottommost plots are classical multidimensional scaling (MDS) plots depicting the separation of individual samples in the indicated clusters. C) Microbial community diversity (Shannon) is reduced as the relative abundance of *P. aeruginosa* (Cluster 1 samples in red and X axis) or *Staphylococcus* spp. (Cluster 2 samples in green) begin to dominate. Individual samples that were clustered in Figure 2A are plotted by the relative abundance of *P. aeruginosa* (x-axis) and their Shannon diversity index (y- axis), then colored by their cluster membership. Samples in Cluster 3 (blue) tended to have higher Shannon diversity values than those in Cluster 1 or 2. D) Relative abundance of *Staphylococcus* spp. overlaid onto the same plot as Figure 2C, showing that Shannon diversity decreases as the relative abundance of this taxon increases.

### Pseudomonas spp. and Staphylococcus spp. drive community structure and low diversity

To further interrogate the drivers of sinus microbial community structure in CF, we performed an unsupervised cluster analysis (Figure 2A). We found that individual sinus samples grouped into three clusters (Supplemental figure 3A), with separation of the two largest clusters driven by the relative abundances of *Pseudomonas* spp. and *Staphylococcus* spp. and the third cluster driven by the relative abundance of a mix of other taxa that were less prevalent in our cohort (Figure 2B). Similarly, these two dominant taxa were also found to unify different groups, with *Pseudomonas* spp. unifying Cluster 1, whereas *Staphylococcus* spp. unified Cluster 2 (Supplemental figure 3B). Consistent with the instability observed in Figure 1, study participants did not tend to belong to solely one cluster. Instead, most participants’ sinus microbiotas frequently switched between clusters over time, depending on the relative abundance of *Pseudomonas* spp. and/or *Staphylococcus* spp. at that time point (Supplemental figure 4).

**Figure 3.**
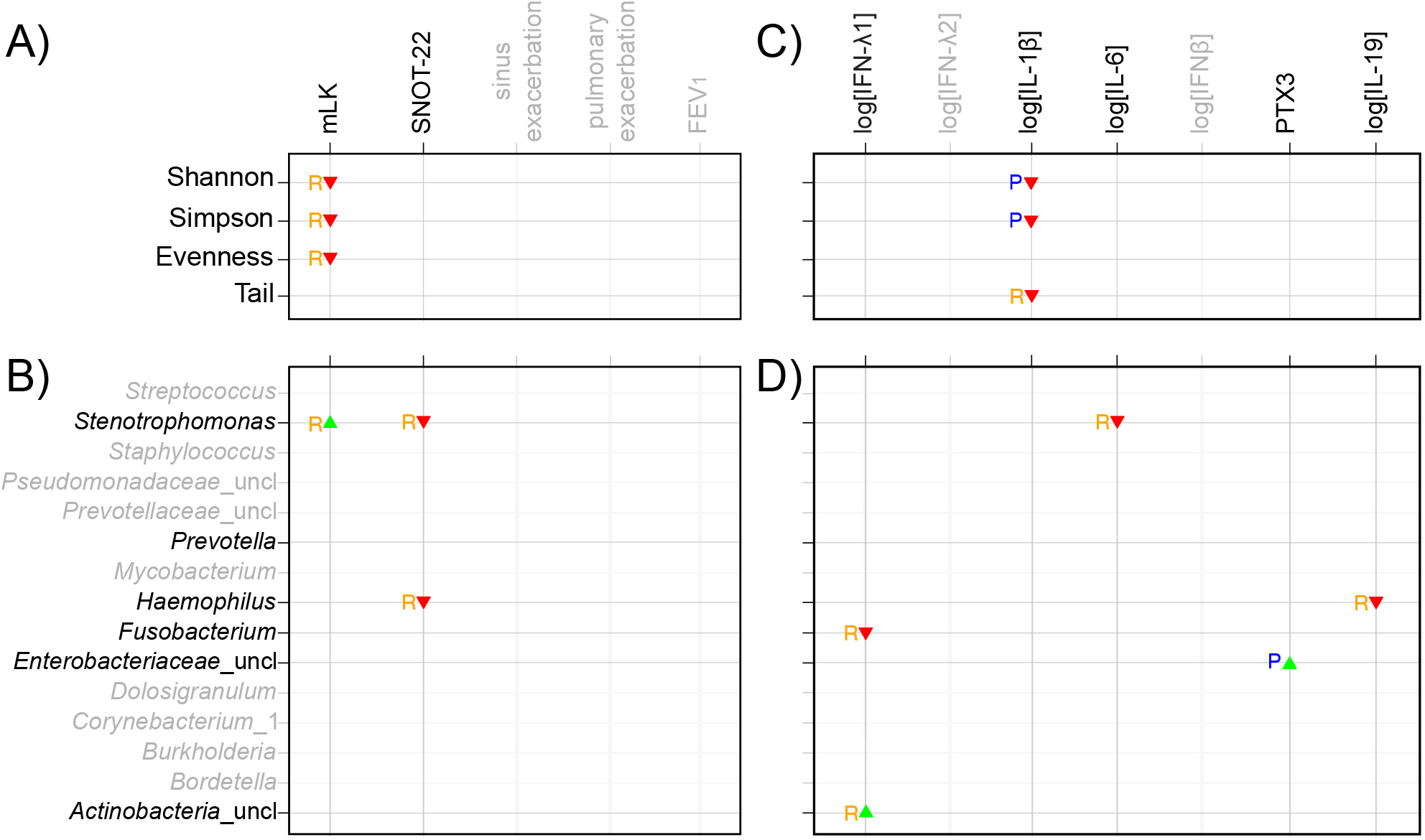
Relationship of microbial community diversity and individual taxa with respiratory disease outcomes and pro- or anti-inflammatory cytokines. Summary matrix depicting statistically significant relationships between clinical outcome variables (left panels: A, B) or cytokines (right panels: C, D) with alpha diversity (top panels: A, C) or the top 15 taxa (lower panels: B, D). The blue P’s represent the associations when the alpha diversity indices or taxa best predict (“P”) the cytokine values or clinical variables. In contrast, the orange R’s depict when the clinical variables or cytokines predict the alpha diversity indices or taxa with greater statistical significance (i.e. when the alpha diversity indices or taxa respond (“R”) to the cytokine values or clinical variables). The green upward-pointing triangles represent positive associations between diversity or taxa and clinical variables or cytokines, whereas the red downward-pointing triangles represent negative associations based on the coefficients of their association. To compute the associations, eight linear regression models were fit. In panel A, 1) diversity indices = covariates + clinical variables and 2) clinical variables = covariates + diversity indices. In panel B, 3.) taxa = covariates + clinical variables and 4) clinical variables = covariates + taxa. In panel C, 5) diversity indices = covariates + cytokines and 6) cytokines = covariates + diversity indices. In panel D, 7.) taxa = covariates + cytokines and 8) cytokines = covariates + taxa. Taxa were represented as additive log ratio transformed abundances. Cytokine concentrations were log transformed, except for pentraxin 3 (PTX3), which was sufficiently normally distributed. Covariates included Patient ID, age, sex, BMI, CFTR mutation, CFRD status, and topical antibiotic usage. Associations included in the predictor/response matrix required at least one of the associations from the models to have an estimated coefficient p-value < 0.025. The coefficients for relationships depicted in each panel are as follows: A) Shannon responds to mLK: -0.138, Simpson responds to mLK: - 0.169, Evenness responds to mLK: -0.138; B) *Stenotrophomonas* responds to mLK: 0.287, *Stenotrophomonas* responds to SNOT-22: -0.0691, *Haemophilus* responds to SNOT-22: -0.0483; B) Shannon predicts log[IL-1*β*]: -2.64, Simpson predicts log[IL-1*β*]: -4.2, Tail responds to log[IL- 1*β*]: -0.233; D) *Stenotrophomonas* responds to log[IL-6]: -1.09, *Haemophilus* responds to log[IL- 19]: -0.287, *Fusobacterium* responds to log[IFN-*λ*1]: -0.342, *Enterobacteriaceae*_uncl predicts PTX3: 341, *Actinobacteria*_uncl responds to log[IFN-*λ*1]: 0.85.

Using the stability classification from Figure 1B, six of the seven individuals with unstable microbiotas switched between clusters in Supplemental figure 5, whereas five of the eight individuals with a stable sinus microbiota exhibited cluster switching. The individual from the unstable group who did not switch clusters (Patient #1 in Supplemental figure 1) stayed within Cluster 3 (cluster driven by a mix of taxa other than *Pseudomonas* spp. or *Staphylococcus* spp.), but their sinus microbiota transitioned from *Streptococcus* spp. to *Burkholderia* spp. dominance, which is why they were grouped among the “Unstable” population despite not switching clusters. In contrast, most of the relatively stable individuals who switched clusters were co-infected with *P. aeruginosa* and *Staphylococcus* spp.; their cluster switching was due to changes in relative abundance of these two taxa that drove them between Clusters 1 (*Pseudomonas*-driven cluster) and 2 (*Staphylococcus*-driven cluster), yet did not lead to a high enough variability in Manhattan distances to categorize them as unstable in Figure 1B because they remained co-infected. Furthermore, the increased relative abundance of *P. aeruginosa* or *Staphylococcus* spp. tracked with decreasing Shannon diversity (Figure 2C,D), suggesting that the low sinus community diversity is attributable to dominance by these two taxa. This reduction in Shannon diversity as the relative abundance of *Pseudomonas* spp. or *Staphylococcus* spp. increased was most apparent in samples from Clusters 1 and 2). Interestingly, CFRD was positively associated with Cluster 1 membership (*Pseudomonas*-driven cluster) and negatively associated with Clusters 2 and 3 (clusters driven by *Staphylococcus* spp. or other taxa) (Figure 2A). In agreement with this clustering analysis, we found that people with CFRD had a higher relative abundance of *Pseudomonas* spp. than those without CFRD (univariate regression coefficient: 45.82, Holm-Bonferroni adjusted p < 0.05). Finally, sinus exacerbation was positively associated with Cluster 2 membership (driven by *Staphylococcus* spp.). These results demonstrate how further clustering sinus microbiotas of people co-infected by *P. aeruginosa* and *Staphylococcus* spp. based on drivers of community structure can reveal relationships with co-morbidities (CFRD) or disease status (sinus exacerbation).

### Sinus microbiotas are highly individualized, but may share a common signature during sinus exacerbation

We used permutational multivariate analysis of variance (PERMANOVA) to determine whether community-level differences in sinus microbiotas track with any characteristics of our study participants or clinical outcomes. We measured sinus disease severity through assessment of symptoms with the validated SNOT-22 questionnaire^14^ and scoring of endoscopic exam findings performed by a clinician (mLK)^15^, and we used FEV1 as an indicator of lung function. We also used sinus or pulmonary exacerbation status at the time of the clinic visit as additional indicators of respiratory disease. We found that the variation in community composition was largely explained by the study participant who contributed the sample (Supplemental table 3; PERMANOVA R^2^ = 0.483, p < 0.001), suggesting that sinus microbiotas are highly individualized. However, whether a person was experiencing a sinus exacerbation at the time of the study visit also explained a small but significant amount of variability (Supplemental table 3; PERMANOVA R^2^ = 0.022, p < 0.05), suggesting that a common signature of disturbance in the CF CRS microbiota may occur during sinus exacerbations.

### Worsened sinus disease is associated with reduced microbial community diversity and changes in Gammaproteobacteria relative abundance

Because the PERMANOVA suggested a distinct microbial community associated with sinus exacerbation and because reduced sputum microbiota diversity is correlated with worse lung function in CF^17–19^, we next asked whether a similar relationship between microbial community diversity and disease occurs in the sinuses during CF CRS. We developed a predictor versus responder test to compare models that use diversity indices or specific taxa as predictors of respiratory disease outcomes against models that use the same indices or taxa as responders to disease outcomes. We found that microbial community diversity (Shannon and Simpson) and evenness decreased in response to increasing mLK score (Figure 3A), suggesting that a more diverse sinus microbiota, not dominated by one or very few taxa, is associated with less severe sinus disease. Examining whether any of the top 15 most abundant taxa were associated with the same disease outcomes (Figure 3B), we found a positive relationship between *Stenotrophomonas* spp. and physician-scored sinus disease (mLK; overall range of 4-16 across study participants’ visits in our study), in which the relative abundance of *Stenotrophomonas* spp. increases in response to increasing mLK score. In contrast, the relative abundance of two taxa, *Stenotrophomonas* spp. and *Haemophilus* spp., decreased in response to worse symptomatic sinus disease (higher SNOT-22 scores). The relative abundance of *Stenotrophomonas* spp. ranged from 0% in some study participants to over 80% in Patient 35’s fourth visit and the relative abundance of *Haemophilus* spp. ranged from 0 up to 17.6% in Patient 18’s second visit (Figure S1). We did not detect statistically significant relationships between these diversity indices or taxa and exacerbation (sinus or pulmonary) or FEV1. Together these data suggest relationships between microbial community diversity, and specifically two Gammaproteobacteria (*Stenotrophomonas* spp. and *Haemophilus* spp.), and upper respiratory disease severity. In the case of the opportunistic pathogen *Stenotrophomonas* spp., the relationship differs depending on whether the outcome of interest is patient- or physician-scored disease.

### Low sinus community diversity and the relative abundance of several taxa are associated with cytokine changes

Both CRS and CF respiratory disease progression are thought to be caused by cycles of infection and inflammation. Therefore, we sought to identify relationships between the sinus inflammatory environment and community diversity, as well as specific taxa. Using the same predictor versus responder approach, we identified a negative relationship between the levels of a pro-inflammatory cytokine and three microbiota diversity indices. Higher Shannon or Simpson diversity predicted decreased IL-1*β* levels (Figure 3C; coefficient [p-value]: Shannon/log[IL-1*β*]: - 2.64 [0.0222], Simpson/log[IL-1*β*]: -4.2 [0.0223]). Similarly, the diversity of low-abundance taxa (Tail statistic) decreased in response to higher levels of IL-1*β* (Figure 3C; -0.233 [0.00058]). Furthermore, the relative abundances of several taxa exhibited relationships with the host cytokine response (Figure 3D). The relative abundances of some taxa decreased in response to increasing levels of cytokines. *Stenotrophomonas* spp. decreased in response to increased levels of pro-inflammatory IL-6 (-1.09 [0.0186], *Haemophilus* spp. decreased in response to anti-inflammatory IL-19 (-0.287 [0.000637], and *Fusobacterium* spp. decreased in response to pro-inflammatory interferon-λ1 (also known as IL-29; -0.342 [0.000653]). The relative abundances of other taxa exhibited positive correlations with some cytokines. Increased relative abundance of unclassified taxa within the family Enterobacteriaceae predicted increased levels of the pro-inflammatory cytokine pentraxin-3 (341.0 [0.0213]), whereas the relative abundance of unclassified taxa within the phylum Actinobacteria increased in response to increasing concentrations of interferon-λ1(0.85 [0.0211]). These findings suggest that sinus inflammation is highest when the microbial community diversity is low.

## Discussion

The unified airway hypothesis suggests that management of upper airway disease (e.g. CRS) could benefit the lower airways, yet the understudied nature of CF sinus disease means evidence-based recommendations for management of CF CRS are currently lacking.^20^ In the present study, we sought to determine how sinus microbial community structure and composition in adults with CF CRS changes across disease and inflammatory states over time. We found that while communities lacked diversity and tended to be dominated by *P. aeruginosa* or *Staphylococcus* spp., many displayed a striking degree of instability over time. Our work also revealed a link between *Staphylococcus* spp. dominance and sinus exacerbation, as well as potential interplay among *P. aeruginosa* dominance in the sinuses, CFRD, and reduced lung function. Pro-inflammatory cytokine responses were associated with decreased sinus microbial community diversity and changes in relative abundance of several taxa. Together these findings shed light on potential host-microbe interactions occurring in the sinuses during CF CRS and have implications for the design of future studies aimed at linking CF CRS phenotypes to disease outcomes, as well as suggest new therapeutic strategies for CF sinus disease.

We identified a striking amount of instability in CF CRS communities over time, even among individuals whose communities were dominated by *P. aeruginosa* or *Staphylococcus* spp. Such instability has been implicated with worse lung function in a meta-analysis of CF sputum microbiotas from several cohorts^21^ and, similarly, in non-CF CRS and diseases involving other microbiome-mucosal interfaces, such as in the gut.^22, 23^ Whether and how the sinus community instability observed in our study is linked to overall respiratory disease progression warrants future investigation, but some parallels can be drawn to existing ecological models of CF lung disease. For example, the climax-attack model of CF pulmonary exacerbation is rooted in the concept of unstable respiratory communities in which the presence/absence or relative abundance of taxa associated with “attack” (community associated with exacerbation, inflammation, and tissue damage) or “climax” (community dominating during periods of clinical stability) communities cycle over time, leading to repeated periods of inflammation, pulmonary exacerbation, and progressive tissue damage.^24^ It is possible that the most unstable communities in our study could be cycling between attack and climax communities, similarly driving sinus disease progression. These communities may also be shifting in response to changing antimicrobial treatments. Another explanation for this instability could be related to the infection site biogeography, specifically the size and structure of bacterial aggregates present in the paranasal sinuses. Using an advanced imaging technique called MiPACT-HCR (<u>m</u>icrobial identification after <u>pa</u>ssive <u>c</u>larity technique and <u>h</u>ybridization <u>c</u>hair reaction)^25^, we recently discovered that the size and structure of *P. aeruginosa* populations varied drastically in adults with CF CRS who were also members of the present cohort, and these varying population sizes impacted ongoing genome evolution and adaptation.^10^ In the present study, we imaged two additional sinus microbial populations (including one that contained *S. aureus*) and observed small, sparse aggregates of bacteria in both participants. Therefore, another explanation for the apparent instability could be due to community compositional differences among isolated aggregates in a structured sinus environment where different taxa may occupy distinct niches. Notably, our examination of microbial community instability is limited by the fact that our microbiota analyses are compositional and not quantitative. Future studies examining the apparent instability of CF CRS microbiotas should endeavor to quantify changes in the absolute abundance of bacteria present at the sampling site.^26, 27^ A future longitudinal study would also benefit from more frequent sample collection with regularly spaced intervals of time between samples to better assess microbial community stability or instability. More broadly speaking, CF respiratory disease is chronic and progressive, with cycles of mucus obstruction, infection, and inflammation driving worsening of respiratory health over an individual’s lifetime.^28^ Predictors of exacerbation or biomarkers of disease progression would greatly inform clinical care, but are currently lacking. The instability identified in our study highlights a potential hurdle for cross-sectional studies that aim to link relative abundance of taxa to outcome measures that develop over longer periods of time (rather than outcomes of acute phenomena), and this limitation should be taken into consideration for future study design.

Our findings share similarities and build upon prior microbiota studies of CF CRS and CF sputum examining diversity and taxonomic drivers of community structure. Consistent with previous adult CF CRS and sputum studies, microbial community diversity was low^4, 5, 17, 29^, and communities were frequently dominated by *P. aeruginosa* or Staphylococci.^4–6, 32–35^ Lucas *et al*. recently showed that low diversity was associated with dominance by *Pseudomonas* spp. and was not significantly associated with clinical factors examined^33^, whereas we found that both *P. aeruginosa* and *Staphylococcus* spp. can drive low community diversity and that low diversity is associated with worse sinus disease as measured by mLK score. Additionally, a recent longitudinal study of four adults with CF CRS reported marked stability in sinus communities over time^35^, whereas approximately half of our cohort exhibited instability. These inconsistencies between studies could be due to differences in study design (e.g. cohort sizes, cross-sectional versus longitudinal), methodological differences in sequencing or analysis approaches, or potentially differences in the cohorts themselves, all of which were recruited at single, distinct study centers. While our 16S amplicon sequencing approach did not detect large taxonomic changes in the group of individuals classified as relatively stable, it is important to note a limitation of this approach. We are unable to determine whether the metabolic activity, for example, of the consortia of stable taxa shift over time in ways that would achieve functionally similar changes as the taxonomic shifts in the relatively “unstable” individuals, as has previously been described in the oral cavity.^36^ On the other hand, taxonomic drivers of the three microbial community clusters identified in our study share similarities with the five clusters recently identified by Hampton *et al*. in a meta-analysis of CF sputum microbiomes from multiple adult cohorts.^21^ We observed a link between cluster membership and sinus disease exacerbation, whereas Hampton *et al*. linked sputum cluster membership to variability in lung function. Specifically, we found that sinus exacerbation was associated with having a microbial community driven by *Staphylococcus* spp. (Cluster 2 membership). This result is consistent with a clinical study of nasal lavages from CF adults which found that colonization of the upper airway by *S. aureus* (as determined by clinical lab culture) was associated with increased levels of several pro-inflammatory cytokines, whereas no association was detected for *P. aeruginosa*.^37^ It is also consistent with the prior publication from our research group on clinical indicators of sinus disease, which found a link between sinus *S. aureus* colonization (as determined by clinical lab culture) and worsening sinus disease.^13^ Together, these studies reveal that beyond resembling each other taxonomically, CF URT and LRT microbial communities also share similar drivers of their structure, some of which can be linked to sinus disease.

CRS is an inflammatory disease and we found several examples of how cytokine signaling in the sinuses could shape CF CRS microbial communities. Multiple lines of evidence link elevated levels of the pro-inflammatory cytokine IL-1*β* to sinus disease. Polymorphisms in the IL-1 receptor antagonist gene (IL1RN) are associated with CRS, and in CF, elevated IL-1*β* is associated with the presence of nasal polyps.^38–41^ We found that lower microbial community diversity was associated with higher levels of IL-1*β* and with worse endoscopic appearance of the sinuses (mLK score). Our findings suggest IL-1*β* signaling in the sinuses as a therapeutic target to control sinus inflammation and potentially restore microbial community diversity. Considering a recombinant IL-1*β* receptor antagonist is already available to treat rheumatoid arthritis and other inflammatory diseases, this finding warrants further investigation.^42^ These cytokine interactions suggest ways that the inflammatory environment of the sinuses could reinforce a lack of diversity and dominance by more abundant taxa, including *P. aeruginosa* and *Staphylococcus* spp.

The CF lung environment displays a high degree of interplay between host-microbe and microbe-microbe interactions that impact bacterial behaviors such as expression of virulence factors and response to antimicrobials.^43^ The strongest signature of microbe-microbe co-occurrence in our study was a positive correlation between *Corynebacterium* spp. and *Dolosigranulum* spp., which has previously been observed in the upper respiratory tract of healthy individuals and associated with relative stability of the microbiota.^44, 45^ We interpret these findings to suggest that CF CRS sinuses continue to harbor a commensal subpopulation displaying interactions seen outside of the context of CF.^44, 46, 47^ We also observed a negative correlation between the relative abundance of *P. aeruginosa* and *Burkholderia* spp. A type VI secretion-mediated mechanism of antagonism between these two organisms was recently demonstrated to evolve among CF sputum isolates, further suggesting similarities between polymicrobial interactions in the URT and LRT.^48^ Finally, we observed a correlation between CFRD and reduced lung function, as was previously reported^13^. Adding to this finding, here we identified a relationship between CFRD and sinus communities driven by *P. aeruginosa* (many of which also contained *Staphylococcus* spp.), which was reminiscent of a report by Limoli *et al*. that LRT co-infection by *P. aeruginosa* and *S. aureus* is common among people with CFRD.^49^ These parallels between our findings and previous reports from studies of the LRT in CF further underscore the relevance of the URT to overall CF respiratory disease dynamics. To further examine CF sinus disease in the context of the unified airway hypothesis, comparison of microbiotas from paired sinus and sputum samples collected longitudinally is needed to examine how these two populations relate to each other and change over time, including during disease exacerbations.

In the United States, the CF Foundation is currently developing guidelines for the management of CF CRS and our study advances the growing body of literature establishing the sinuses as an important site of chronic infection along the respiratory tract of people with CF. More work is needed in a larger cohort to understand the causes and consequences of sinus microbial community instability in adults with CF CRS, and how community structure relates to the inflammatory environment of the sinuses. Furthermore, the CF community has entered the era of highly effective modulator therapy (HEMT), in which widespread use of HEMT is changing CF disease in unprecedented ways that will require complementary changes in how CF is managed for many people^50^. Initiation of HEMT early in life may delay respiratory disease progression and it is possible that management of sinus disease could offer an additional opportunity to prevent or delay LRT disease, as well as to relieve the symptomatic quality of life burden for people with CF. As therapeutic options for CF are expanded and life expectancy extended, a comprehensive understanding of the ecological and evolutionary drivers of CF sinus disease holds promise for more rational interventions and treatment of chronic respiratory infections in people with CF.

## Methods

### Study design, participants, sinus sampling, and clinical evaluation

We performed a prospective, longitudinal study of 33 CF adults with symptomatic CRS and prior functional endoscopic sinus surgery (FESS) following an IRB-approved protocol (STUDY19100149) between February 2015 and August 2017.^13^ Participants were treated in a CF-focused otolaryngology clinic at the University of Pittsburgh. During quarterly clinic visits and unscheduled clinic visits, at least two sinus swabs were collected endoscopically for 16S rRNA gene amplicon sequencing (dry flock swab; Puritan Medical Products, Guilford, Maine) and bacterial culturing (flocked swab with liquid Amies medium; Copan Diagnostics, Inc. Murrieta, CA). Samples were collected from the frontal, maxillary, or ethmoid sinuses (Supplemental Table 4). The swab for bacterial culturing was stored on wet ice and cultured within 4 hours of sampling. Sinus wash was collected for cytokine analysis by flushing 5mL of sterile saline into the sinus cavity and collecting endoscopically with a sterile trap. Sinus washes and dry swabs were stored at -80°C. Of the 33 participants enrolled in the study and for whom sinus samples had been collected, we sequenced microbiota samples on at least two different study visits from 18 people (i.e. longitudinal samples) and sequenced a single cross-sectional sample from 9 people, for a total cohort size of 27 people (Table 1). Patient demographics, clinical characteristics including the criteria for disease outcome variables “sinus exacerbation” and “pulmonary exacerbation”, and medication use was previously described for the full 33 person cohort and the same definitions were used in the present study.^13^ Briefly, a sinus disease exacerbation was defined as an unscheduled visit to the sinus clinic (i.e. a visit that was outside of regular study visits) and/or if the study participant reported an acute increase in symptom severity. A pulmonary exacerbation occurred if at least two of the following three occurred within four weeks of a study visit: (1) a greater than 10% drop in percent predicted FEV1, (2) institution of a new course of systemic antibiotics by the pulmonary team, and (3) documentation by the treating pulmonologist that the study participant was experiencing a pulmonary exacerbation in the medical record. Supplemental table 4 contains the study’s clinical metadata and Supplemental table 5 is a codebook describing each variable.

### DNA extraction and 16S rRNA gene amplicon sequencing

DNA extraction was performed using the Qiagen DNeasy Powersoil Kit (Qiagen Cat#12888, Germantown, MD) and processed following the manufacturer’s protocol. Reagent blanks were included as negative controls and cells from a microbial community of known composition were included as positive controls (ZymoBiomics Microbial Community Standards; Zymo Research, Irvine, CA). The V4 region of the 16S rRNA gene was amplified from approximately 5 ng of extracted DNA in 25μl reactions using Q5 HS High-Fidelity polymerase (New England BioLabs, Ipswich, MA) with inline bare primer design as previously described.^51^ The following V4-specific primers were used: 515f 5’-GTGCCAGCMGCCGCGGTAA-3’ and 806r 5’- GGACTACHVGGGTWTCTAAT-3’. Cycle conditions were 98°C for 30 seconds, followed by 30 cycles of 98°C for 10 seconds, 57°C for 30 seconds, and 72°C for 30 seconds, then a final extension step of 72°C for 2 minutes. We used two-sided AMPure XP bead purification at 0.8:1 (left-side) and 0.61:1 (right-side) ratios to remove small and large fragments, respectively. Eluted DNA was quantified on a Qubit fluorimeter (Life Technologies, Grand Island, NY). Samples were pooled on ice by combining 40ng of each purified band. For negative controls and poorly performing samples, 20μl of each sample was used. The sample pool was purified with the MinElute PCR purification kit (Qiagen, Germantown, MD). The final sample pool underwent two more purifications: AMPure XP beads at a ratio of 0.8:1 to remove primer dimers and a final cleanup using the Purelink PCR Purification Kit (Life Technologies Cat #K310001; Grand Island, NY). The purified pool was quantified in triplicate with a Qubit fluorimeter prior to sequencing.

Amplicons of the V4 region were sequenced on a MiSeq (Illumina, San Diego, CA) using paired-end 2 x 250 reads, deconvolved, and quality checked by dust low complexity filtering, quality value (QV) trimming, and trimming of primers used for 16S rRNA gene amplification by the University of Pittsburgh’s Center for Medicine and the Microbiome (CMM) using the scripts fastq_quality_trimmer and fastq_quality_filter from Hannon’s Cold Spring Harbor Laboratory’s FASTAX-Toolkit (http://hannonlab.cshl.edu/fastx_toolkit/). Reads were trimmed until the QV was 30 or higher. Trimmed reads shorter than 75bp or those with less than 95% of the bases above a QV of 30 were discarded. Forward and reversed paired reads were merged with a minimum required overlap of 25 bp, proportion overlap mismatch > 0.2 bp, maximum N’s allowed = 4, and a read length minimum of 125 bp. Reads were taxonomically classified with Mothur version 1.39.1^16^, using Ribosomal Database Project (RDP v123) reference sequences.^52^ Environmental controls and extraction kit controls, along with *E. coli* and mock community (ZymoBIOMICS Microbial Community DNA Standard) positive controls, were sequenced alongside clinical specimens to monitor for contamination and technical performance during the extraction and sequencing process.

### Verification of Pseudomonadaceae_uncl taxon as P. aeruginosa

For every study visit, a sinus swab was streaked onto *Pseudomonas* isolation agar (PIA) and incubated at 37°C for 48 hours. Genomic DNA from representative isolate(s) for each study participant was extracted using a QIAGEN DNeasy Blood & Tissue kit (Qiagen, Hilden, Germany) and the 16S rRNA gene was amplified using the primers 63f and 1387r.^53^ Amplicons were purified enzymatically with ExoSAP-It (Applied Biosystems, Waltham, MA) prior to Sanger sequencing (Eurofins Genomics, Louisville, KY) to confirm their species identity as *P. aeruginosa*. We did not detect Pseudomonads other than *P. aeruginosa*. Furthermore, whole genomes were previously sequenced for all *P. aeruginosa* isolates collected from six study participants (Patients 9, 24, 32, 33, 41, and 52 in Figure S1). We did not detect non-*P. aeruginosa* Pseudomonads among whole genome sequenced isolates.

### FISH imaging of explanted obstructive sinus material

When clinically indicated, obstructive sinus material was surgically removed from two study participants following sinonasal endoscopy and immediately fixed in 10% phosphate-buffered formalin (Fisher Scientific). Fixed samples were then rinsed and embedded for freezing in O.C.T. compound (Tissue-Plus, Fisher HealthCare). Cryoprotected samples were sectioned at 10µm on a Microm HM505E Cryostat Microtome (Microm International, Waldorf, Germany) and immobilized on poly-L-lysine coated slides. In preparation for staining, slides were removed from the freezer and thawed at room temperature. Samples were permeabilized by incubating with lysozyme (10 mg/mL in 0.1M Tris-HCL and 0.05 M EDTA) at 37°C for 3 hours. The lysozyme solution was removed, and the samples were rinsed briefly with sterile RNase/DNase-free water. The samples were then dehydrated with increasing concentrations of ethanol (50%, 80% and 100%) for 3 minutes per treatment as previously described^54^ and then air dried at room temperature. FISH was performed using oligonucleotide probes directed toward 16S rRNA sequences specific to Eubacteria (Eub338; 5’-GCT GCC TCC CGT AGG AGT-3’)^55^, *S. aureus* (Sau16S69; 5’-GAA GCA AGC TTC TCG TCC G-3’)^56^, or *P. aeruginosa* (PsaerA; 5’-GGT AAC CGT CCC CCT TGC-3’).^54^ Probes were synthesized by IDT (Coralville, IA) and 5’ labeled with the cyanine dye Cy3 (PsaerA and Sau16S69) or Cy5 (Eub338). Samples were incubated in hybridization buffer (0.9 M NaCl, 20 mM Tris-HCl [pH 7.6], 0.01% sodium dodecyl sulfate, 30% formamide) with desired probe combinations for 1h at 46°C. Samples were then washed with pre- warmed washing buffer (20 mM Tris-HCl [pH 7.6], 0.01% sodium dodecyl sulfate, 112 mM NaCl) and incubated in washing buffer for 15 minutes at 48°C. Slides were then rinsed with sterile water, and the general DNA stain Hoechst trihydrochloride trihydrate was applied (1.0 µg/mL in PBS) for 10 minutes on ice. Slides were rinsed again with sterile water and left to air dry in a vertical position protected from light. When dry, samples were mounted with ProLong Gold antifade reagent (Life Technologies) for microscopy. Microscopy was performed on an Olympus FluoView FV1000 inverted confocal microscope using a 60X oil objective.

### Cytokine panels

Cytokine levels in 54 sinus washes (stored frozen at -80°C) from 22 people were quantified using a Bio-Plex Pro™ Human Inflammation 24-Plex Panel or a Bio-Plex Pro™ Human Th17 Cytokine Panel 15-Plex Panel (Bio-Rad, Hercules, CA, USA). Cytokines were omitted from further analyses if their concentration was close to the lower limit of detection in most samples, based on manufacturer-specified values and examination of the 5pl standard curves produced by the Bio- Plex Pro™ software. Seven cytokines were included in the predictor/responder analyses in Figure 3. All cytokine concentrations were log-transformed except pentraxin 3 (PTX3), which was sufficiently normally distributed according to the Shapiro-Wilks test, without additional log-transformation.

### Statistical analyses

Statistical analyses were performed with GraphPad Prism version 9.1.2 or in RStudio version 1.1.456, and significance determined at *α* = 0.05 unless otherwise specified. Relative abundances of taxa were transformed using the additive log ratio (ALR) transformation.^57, 58^ Alpha diversity values were calculated using the *diversity* function in the R package vegan v2.5-3. The clustering analyses in Figure 2A, B and Figure S3 were performed with vegan and the R packages permute and lattice. The Manhattan distance between each sample was computed, and samples were hierarchically clustered based on the Ward’s minimum variance method. The taxon/taxa driving cluster formation was determined by calculating R^2^ ratios (sum of squares between clusters divided by sum of squares total) from ANOVA. In Figure 2A, cluster multinomial log linear models were fit for covariates and clinical outcome variables to determine relationships between any of these variables with 16S microbiota profile cluster(s). The PERMANOVA in Supplemental table 3 was performed with the *adonis* function in vegan (N = 88 samples; 12000 permutations). Linear models in the predictor/responder analyses in Figure 3 were calculated as previously described.^59^ Briefly, for both the sinus disease outcome and cytokine analyses, the following covariates were included Patient ID (to control for repeated measures within patients), age, sex, BMI on enrollment, CFTR mutation, CFRD diagnosis, and current topical antibiotic use. A p-value < 0.025 in both the predictor and responder versions of the versions of the model (i.e., alpha = 0.05 for two simultaneous tests) was required for relationships to be summarized in Figure 3. Taxon co-occurrences in Figure S2 were determined for the top 15 taxa by Pearson correlation across all samples (N=101) and significance was determined after Holm-Bonferroni correction (p < 0.05). The Rs95 values included with taxon prevalence in Supplemental table 2 provide estimates that account for unequal sequencing depth using the binomial distribution. For example, in a sample with a read depth of 3000, if the abundance of the taxon was 0.001, then according to the binomial distribution, the probability of not detecting this taxon (0 reads) is 0.0497, if the sample was re-sequenced to the same depth. Therefore, at least 1 read will be associated with this taxon in that sample, with a probability of 1-0.0497 = 0.9505 or > 95% of the time.

### Data sharing

All V4 amplicon sequencing reads were deposited in NCBI’s SRA under BioProject PRJNA750353. Further information and requests for resources and reagents should be directed to and will be fulfilled by the lead contact, Dr. Jennifer Bomberger.

## Acknowledgements

We thank the people with CF who participated in our study and contributed to this research. We also thank Bill Goins and Mingdi Zhang for assistance with cryosectioning of clinical samples. This work was supported by a Cystic Fibrosis Foundation (CFF) Carol Basbaum Memorial Research Fellowship (ARMBRU19F0) and National Institutes of Health (NIH) T32HL129949 to CRA, NIH T32AI049820 and CFF MELVIN15F0 to JAM, CFF ZEMKE16A0 and NIH 5K23HL131930 to ACZ, NIH R01HL138630 to AM, NIH R21HL143091 to BAM, University of Pittsburgh CTSI Pilot Program, NIH NCATS UL1 TR0000005, NIH R61HL137077 and GILEAD Investigator Sponsored Research Award to SEL and JMB, and NIH R01HL123771, P30DK072506, CFF BOMBER14G0, and CFF RDP BOMBER19R0 to JMB.

## Supplemental figure legends

**Figure S1.**
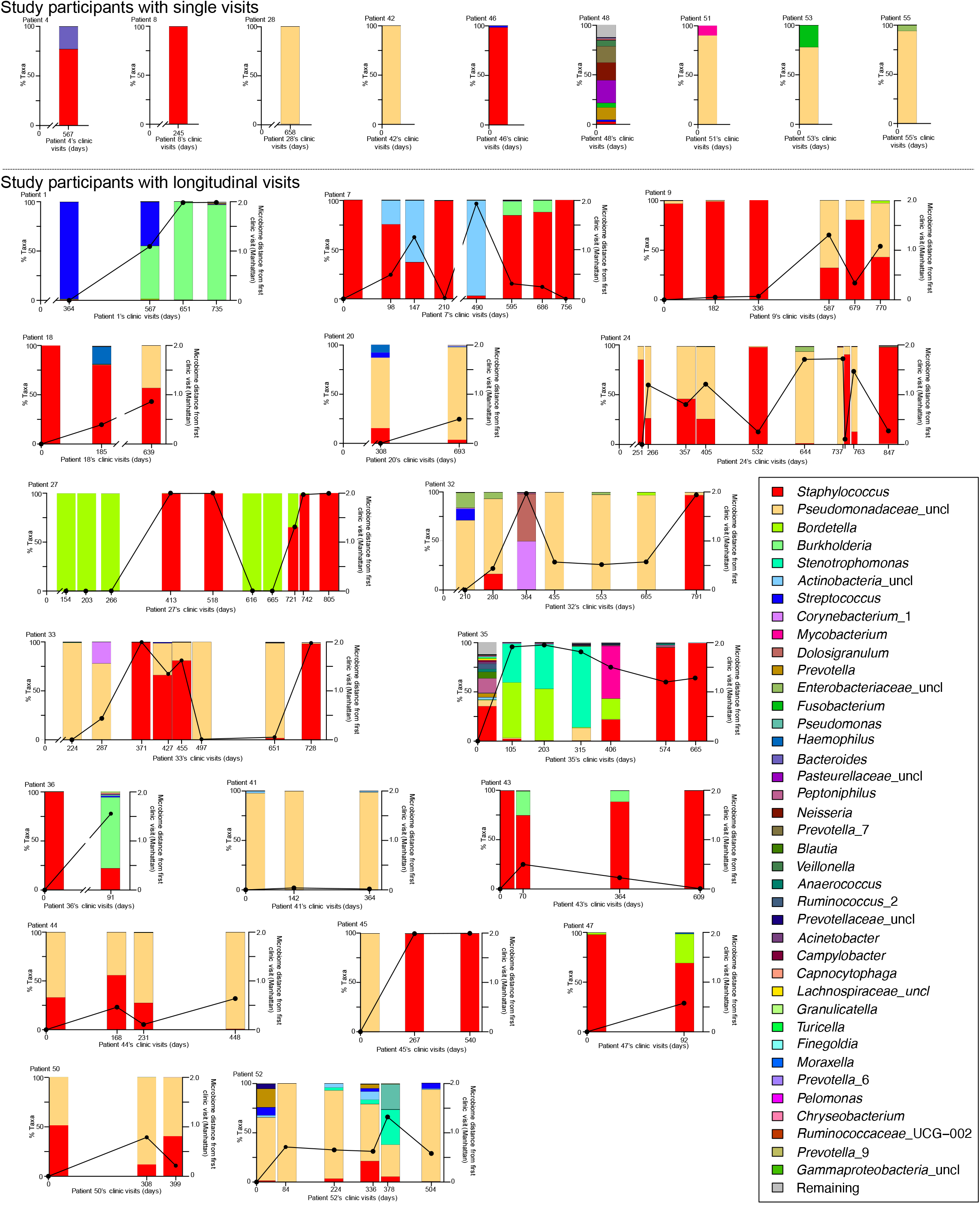
CF CRS microbial communities can be unstable, with many individuals switching between *Staphylococcus* spp. and *P. aeruginosa* at the greatest levels of relative abundance over time. Taxa bar plots depicting the percent of each taxon measured at clinic visit dates for each of the 27 study participants for whom we sequenced one microbiota sample (top) or longitudinal (bottom) samples. Colors representing each taxa can be matched by the legend on the right. *Staphylococcus* spp. is depicted in red, *P. aeruginosa* is represented by *Pseudomonadaceae* in peach and *Pseudomonas* spp. in dark teal. Values indicated beneath each stacked bar plot signify the days since study enrollment. Black lines were drawn over the individual bar plots to indicate the degree of microbiota dissimilarity relative to the first time point. The units of dissimilarity were measured with the Manhattan distance and are annotated on the y-axis on the right.

**Figure S2.**
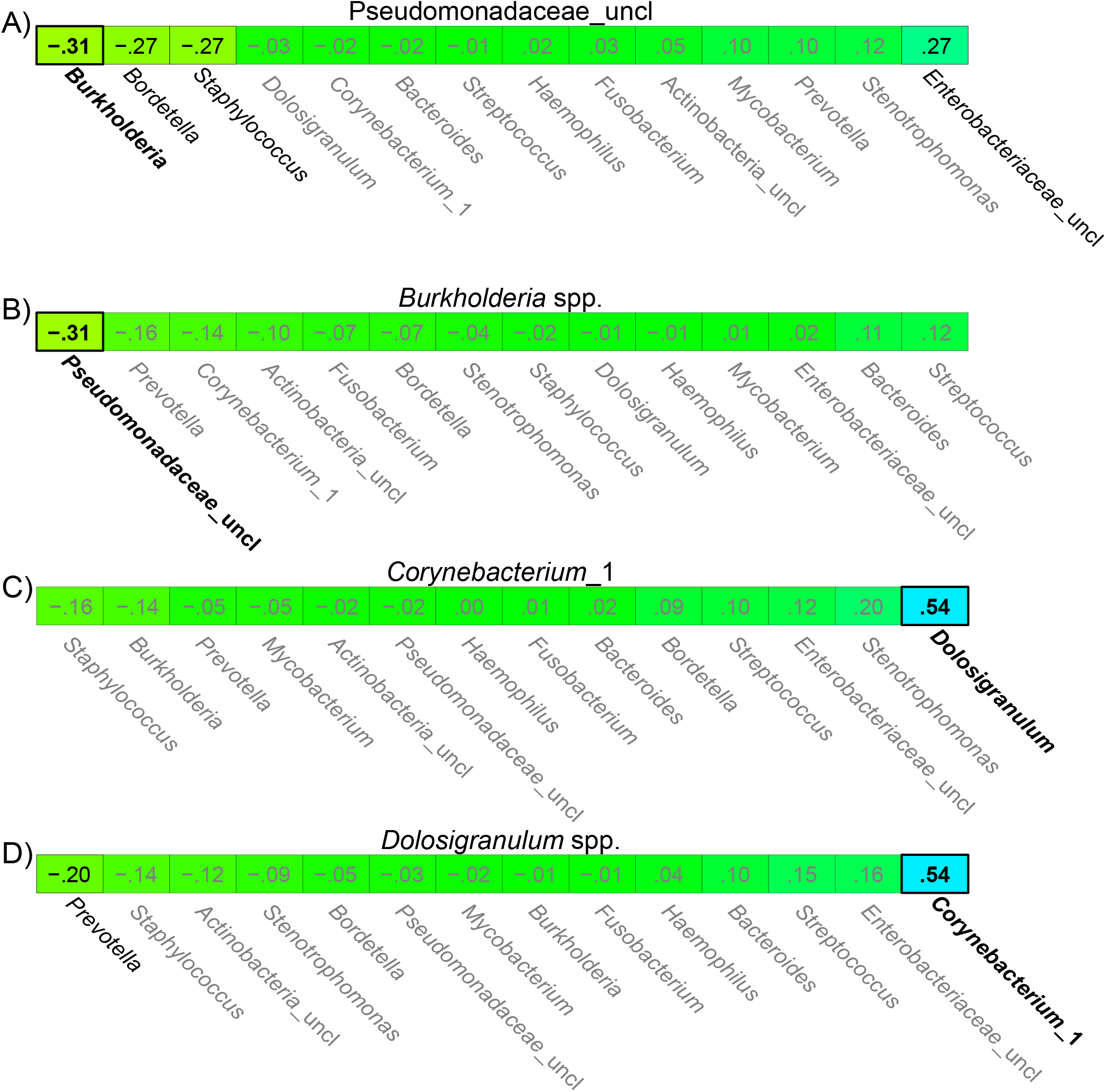
Commensal taxa *Corynebacterium* spp. and *Dolosigranulum* spp. co-occur, whereas opportunistic pathogens *P. aeruginosa* and *Burkholderia* spp. display antagonism. Depicted in bold text and with a thick black box are the coefficients for relationships among the top 15 taxa that were statistically significant after controlling for multiple comparisons (Holm-Bonferroni adjusted p-value <0.05). In non-bold black text are individual associations that were statistically significant prior to correcting for multiple hypothesis testing (p < 0.05), but not after (Holm-Bonferroni adjusted p-value >0.05).

**Figure S3.**
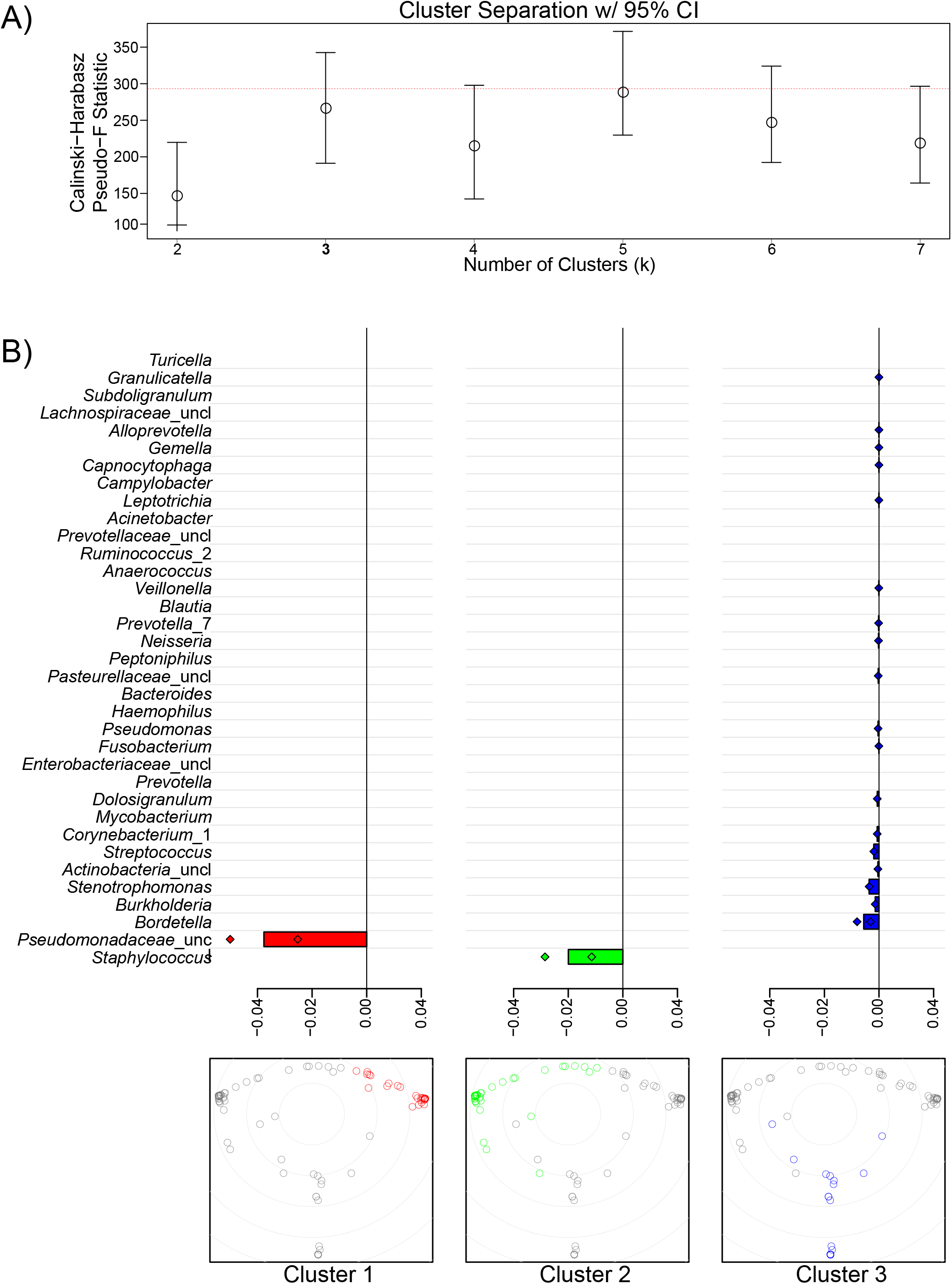
Characterization of cluster assignment in. **Figure 2. A)** The Calinski-Harabasz Pseudo-F statistic was calculated across all cluster cuts (k) to determine the optimal numbers of cluster cuts to use in Figure 2AB. Although at k = 5 cluster cuts, the clusters had the greatest inter-cluster separation, the cuts from k = 3 to 7 were not statistically significantly different from each other. Ultimately, the cluster cut at k = 3 was chosen because deeper cuts at k = 4 or k = 5 would have yielded cluster sizes too small to find statistically significant associations with the clinical or cytokine data. **B)** To determine which taxa influence the differentiation of clusters from one another (in Figure 2AB), taxa were iteratively evaluated for their contribution to pair-wise cluster separation by comparing the coefficient of determination (R^2^) with (full) and without (reduced) the taxon of interest. The x-axis annotates this calculated metric: log(R^2^ reduced / R^2^ full). If excluding a taxon (reduced model) increases the separation between two clusters relative to its inclusion (full model), then it was an important clustering influencer, and the log ratio would be <1. Log ratios greater than 1 indicate that the taxon added more noise (within cluster variance), thus reducing between cluster separation.

**Figure S4.**
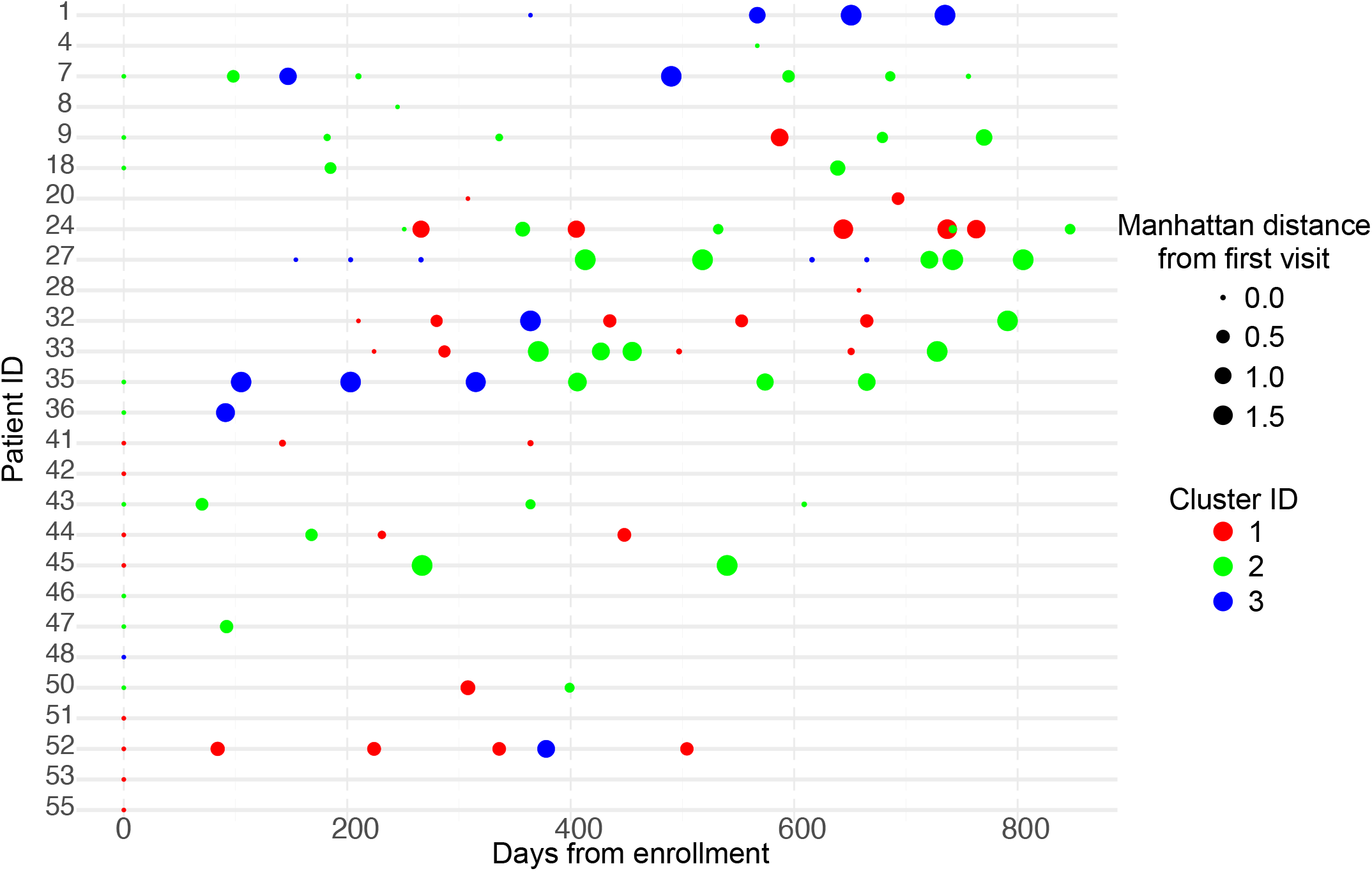
Twelve of eighteen individuals with longitudinal microbiota samples switch between cluster membership over time. Cluster membership is colored as in Figure 2. Cluster 1 (*P. aeruginosa*) = red, Cluster 2 (*Staphylococcus* spp.) = green, Cluster 3 (other taxa) = blue. The size of each dot is proportional to the Manhattan distance at each timepoint from the first sequenced sample.

## Supplemental table legends

**Supplemental table 1.** Diversity indices for each sequenced microbiota sample. The sample ID numbers listed correspond to the sample ID numbers in the metadata file (Supplemental table 4).

| Sample ID | Shannon | Simpson | Evenness | Tail | Richness |
| --- | --- | --- | --- | --- | --- |
| 0104.01.01.SINUS | 0.10589483 | 0.03397046 | 0.05092465 | 0.287282 | 8 |
| 0104.01.02.SINUS | 0.8179829 | 0.51611268 | 0.27780603 | 1.23346473 | 19 |
| 0104.01.03.SINUS | 0.05154394 | 0.01166583 | 0.01750552 | 0.62712527 | 19 |
| 0104.01.04.SINUS | 0.23749582 | 0.06885968 | 0.08769993 | 1.23729498 | 15 |
| 0104.04.01.SINUS | 0.55157206 | 0.35447306 | 0.34271099 | 0.48696288 | 5 |
| 0104.07.01.SINUS | 0.5631518 | 0.37064178 | 0.16249126 | 0.57228765 | 32 |
| 0104.07.02.SINUS | 0.66782445 | 0.46875164 | 0.34319388 | 0.61981386 | 7 |
| 0104.07.03.SINUS | 0.07910591 | 0.01762764 | 0.0211645 | 1.57350577 | 42 |
| 0104.07.04.SINUS | 0.15674374 | 0.06603898 | 0.1426743 | 0.20127525 | 3 |
| 0104.07.05.SINUS | 0.50714676 | 0.26517697 | 0.15957777 | 0.98347694 | 24 |
| 0104.07.06.SINUS | 0.40024958 | 0.21522894 | 0.14779991 | 0.59603557 | 15 |
| 0104.07.07.SINUS | 0.0121703 | 0.00264745 | 0.00679237 | 0.11119694 | 6 |
| 0104.07.08.SINUS | 0.00778681 | 0.00148121 | 0.00324735 | 0.16889304 | 11 |
| 0104.08.01.SINUS | 0.00432632 | 0.00088028 | 0.00312078 | 0.04532762 | 4 |
| 0104.09.01.SINUS | 0.08402852 | 0.02486021 | 0.02470557 | 0.57216535 | 30 |
| 0104.09.02.SINUS | 0.00142403 | 0.00028937 | 0.00205443 | 0.01202944 | 2 |
| 0104.09.03.SINUS | 0.65588247 | 0.43865674 | 0.36605497 | 0.61119325 | 6 |
| 0104.09.04.SINUS | 0.51529151 | 0.31923503 | 0.23451927 | 0.47379282 | 9 |
| 0104.09.05.SINUS | 0.78229205 | 0.5148748 | 0.4366055 | 0.72857792 | 6 |
| 0104.09.06.SINUS | 0.19234726 | 0.06978859 | 0.0605236 | 0.93817563 | 24 |
| 0104.18.01.SINUS | 0.62057285 | 0.32374802 | 0.16939058 | 1.92072164 | 39 |
| 0104.18.02.SINUS | 0.68887458 | 0.49111112 | 0.49691797 | 0.65939088 | 4 |
| 0104.18.03.SINUS | 0.00836935 | 0.00174383 | 0.00467103 | 0.08647293 | 6 |
| 0104.20.01.SINUS | 0.89683958 | 0.46001571 | 0.40816928 | 0.98816278 | 9 |
| 0104.20.04.SINUS | 0.3885309 | 0.13060842 | 0.09652098 | 2.68572267 | 56 |
| 0104.24.01.SINUS | 0.4094465 | 0.23690178 | 0.17782036 | 0.41456201 | 10 |
| 0104.24.02.SINUS | 0.58119984 | 0.39109617 | 0.5290309 | 0.5167745 | 3 |
| 0104.24.03.SINUS | 0.7110311 | 0.50035517 | 0.44178846 | 0.69425102 | 5 |
| 0104.24.04.SINUS | 0.63897899 | 0.39483391 | 0.19850998 | 1.16587182 | 25 |
| 0104.24.05.SINUS | 0.05833955 | 0.02057431 | 0.05310295 | 0.10332065 | 3 |
| 0104.24.07.SINUS | 0.30522775 | 0.12649838 | 0.12283268 | 0.5061858 | 12 |
| 0104.24.08.SINUS | 0.29419649 | 0.15405914 | 0.21221791 | 0.29673778 | 4 |
| 0104.24.09.SINUS | 0.42659598 | 0.23561198 | 0.26505899 | 0.41326579 | 5 |
| 0104.24.10.SINUS | 0.05857163 | 0.01761035 | 0.0363926 | 0.1517269 | 5 |
| 0104.24.12.SINUS | 0.10335155 | 0.03279684 | 0.04703732 | 0.28326927 | 9 |
| 0104.27.01.SINUS | 0.01002628 | 0.00181389 | 0.00289297 | 0.40264721 | 32 |
| 0104.27.02.SINUS | 0.0079806 | 0.00161679 | 0.00410122 | 0.10310041 | 7 |
| 0104.27.03.SINUS | 0.02148301 | 0.00472933 | 0.00932995 | 0.21063452 | 10 |
| 0104.27.04.SINUS | 0.03243851 | 0.00824179 | 0.01305422 | 0.23774389 | 12 |
| 0104.27.05.SINUS | 0.0210052 | 0.00565583 | 0.01515205 | 0.06893748 | 4 |
| 0104.27.06.SINUS | 0.01964533 | 0.00571675 | 0.02834221 | 0.05354057 | 2 |
| 0104.27.07.SINUS | 0.01097545 | 0.00263642 | 0.00791711 | 0.05137333 | 4 |
| 0104.27.08.SINUS | 0.65581038 | 0.45334025 | 0.4074779 | 0.59864025 | 5 |
| 0104.27.09.SINUS | 0.07297444 | 0.02729362 | 0.10527986 | 0.11763635 | 2 |
| 0104.27.10.SINUS | 0.0177899 | 0.00440411 | 0.0128327 | 0.08401063 | 4 |
| 0104.28.01.SINUS | 0.02275625 | 0.00551117 | 0.0116944 | 0.14182516 | 7 |
| 0104.32.01.SINUS | 0.86733327 | 0.45454917 | 0.36170607 | 0.95381009 | 11 |
| 0104.32.02.SINUS | 0.69004526 | 0.37816213 | 0.35461312 | 0.67418798 | 7 |
| 0104.32.03.SINUS | 0.82344377 | 0.52160291 | 0.28489199 | 0.99783591 | 18 |
| 0104.32.04.SINUS | 0.02379887 | 0.00605704 | 0.0122302 | 0.1255229 | 7 |
| 0104.32.05.SINUS | 0.14457184 | 0.05562015 | 0.06278675 | 0.25443824 | 10 |
| 0104.32.06.SINUS | 0.15334763 | 0.06380262 | 0.07880509 | 0.21475188 | 7 |
| 0104.32.07.SINUS | 0.15684413 | 0.0605612 | 0.06811654 | 0.34175494 | 10 |
| 0104.33.01.SINUS | 0.04310884 | 0.01254171 | 0.01491464 | 0.27159448 | 18 |
| 0104.33.02.SINUS | 0.53595587 | 0.34303814 | 0.27542683 | 0.48392336 | 7 |
| 0104.33.03.SINUS | 0.03000626 | 0.00746224 | 0.01365644 | 0.18039856 | 9 |
| 0104.33.04.SINUS | 0.69416067 | 0.45336895 | 0.24500826 | 0.73161372 | 17 |
| 0104.33.05.SINUS | 0.49254924 | 0.30718119 | 0.21391142 | 0.46863302 | 10 |
| 0104.33.06.SINUS | 0.00719502 | 0.00139141 | 0.00280513 | 0.1381955 | 13 |
| 0104.33.07.SINUS | 0.1788408 | 0.05278516 | 0.045953 | 1.98165623 | 49 |
| 0104.33.08.SINUS | 0.08335648 | 0.02719195 | 0.03620126 | 0.25233094 | 10 |
| 0104.35.01.SINUS | 0.89561934 | 0.52979487 | 0.32302639 | 1.00465728 | 16 |
| 0104.35.02.SINUS | 0.85395521 | 0.52355481 | 0.26870382 | 1.25972202 | 24 |
| 0104.35.03.SINUS | 0.75738899 | 0.31301841 | 0.19782187 | 3.56206679 | 46 |
| 0104.35.04.SINUS | 1.24399943 | 0.62473957 | 0.31799389 | 3.13508178 | 50 |
| 0104.35.05.SINUS | 0.3378025 | 0.10282788 | 0.08468384 | 2.96338754 | 54 |
| 0104.35.06.SINUS | 2.63562916 | 0.83780344 | 0.62247588 | 13.0473956 | 69 |
| 0104.35.07.SINUS | 0.06032969 | 0.01531783 | 0.02427845 | 0.38003598 | 12 |
| 0104.36.01.SINUS | 0.03495463 | 0.0098714 | 0.01457721 | 0.17654247 | 11 |
| 0104.36.02.SINUS | 0.83224584 | 0.42967049 | 0.42768976 | 1.14070365 | 7 |
| 0104.41.01.SINUS | 0.10540975 | 0.04214031 | 0.07603706 | 0.15114284 | 4 |
| 0104.41.02.SINUS | 0.01533826 | 0.0039999 | 0.01106422 | 0.05744511 | 4 |
| 0104.41.03.SINUS | 0.06672001 | 0.02447487 | 0.09625663 | 0.11131475 | 2 |
| 0104.42.01.SINUS | 0.01807113 | 0.00442783 | 0.01303556 | 0.09118583 | 4 |
| 0104.43.01.SINUS | 0.02637757 | 0.00614694 | 0.00866394 | 0.3089862 | 21 |
| 0104.43.02.SINUS | 0.59837818 | 0.37762027 | 0.21581931 | 0.67459121 | 16 |
| 0104.43.03.SINUS | 0.36706226 | 0.20307205 | 0.14771672 | 0.42625751 | 12 |
| 0104.43.04.SINUS | 0.03255379 | 0.0070786 | 0.01233538 | 0.45574127 | 14 |
| 0104.44.01.SINUS | 0.64386079 | 0.44264048 | 0.21492601 | 0.659217 | 20 |
| 0104.44.02.SINUS | 0.69408029 | 0.49392019 | 0.31588955 | 0.67550772 | 9 |
| 0104.44.03.SINUS | 0.61347552 | 0.40354821 | 0.21224796 | 0.68952096 | 18 |
| 0104.44.05.SINUS | 0.06140531 | 0.02132493 | 0.04429457 | 0.11321719 | 4 |
| 0104.45.01.SINUS | 0.0081712 | 0.00171266 | 0.00456044 | 0.08352691 | 6 |
| 0104.45.02.SINUS | 0.03262632 | 0.01001701 | 0.02027187 | 0.08118358 | 5 |
| 0104.45.05.SINUS | 0.01537737 | 0.00400778 | 0.01399708 | 0.05487295 | 3 |
| 0104.46.01.SINUS | 0.10567839 | 0.03882913 | 0.03902379 | 0.34435305 | 15 |
| 0104.47.01.SINUS | 0.09856943 | 0.03622011 | 0.04280816 | 0.2160261 | 10 |
| 0104.47.02.SINUS | 0.66867394 | 0.43215508 | 0.37319403 | 0.58890149 | 6 |
| 0104.48.01.SINUS | 2.35363069 | 0.86435954 | 0.63379157 | 5.61677333 | 41 |
| 0104.50.01.SINUS | 0.69408007 | 0.49985878 | 0.6317789 | 0.69914799 | 3 |
| 0104.50.02.SINUS | 0.37886231 | 0.21033836 | 0.19469672 | 0.37760126 | 7 |
| 0104.50.03.SINUS | 0.68887166 | 0.48359093 | 0.38446659 | 0.64854983 | 6 |
| 0104.51.01.SINUS | 0.34527223 | 0.18211599 | 0.13461171 | 0.46116123 | 13 |
| 0104.52.01.SINUS | 1.12138729 | 0.54057118 | 0.40445497 | 1.41079921 | 16 |
| 0104.52.02.SINUS | 0.00520951 | 0.00105428 | 0.00290748 | 0.05420299 | 6 |
| 0104.52.03.SINUS | 0.44870722 | 0.19058439 | 0.19487107 | 0.69406454 | 10 |
| 0104.52.05.SINUS | 1.2623419 | 0.60501169 | 0.39720595 | 1.70145422 | 24 |
| 0104.52.06.SINUS | 1.29989416 | 0.70018226 | 0.66801345 | 1.43817256 | 7 |
| 0104.52.07.SINUS | 0.27742563 | 0.12400344 | 0.20012029 | 0.33000668 | 4 |
| 0104.53.01.SINUS | 0.5450781 | 0.34868089 | 0.28011473 | 0.49035187 | 7 |
| 0104.55.01.SINUS | 0.24599724 | 0.11865297 | 0.1774495 | 0.26771423 | 4 |

**Supplemental table 2.** Top 100 taxa identified in this study. The mean abundance per participant is averaged across all study participants, regardless of whether the taxon was detected. For participants with multiple study visits, the relative abundance of each taxon was first averaged across their visits. Number of participants is a count of the number of study participants with at least one study visit in which that taxon was identified. The prevalence is the percentage of patients with at least one study visit in which that taxon was identified. The Rs95 value is an estimate of mean abundance that takes into account the uneven sequencing depth of samples and is presented as a count of participants and prevalence based on this adjustment. All statistics for taxonnomic abundances are performed on additive log-transformed abundances.

| Rank | Taxa | Mean abundance<br>per participant | Number of<br>participants | Prevalence (%) | Number of<br>participants<br>(Rs95) | Prevalence<br>(Rs95;%) | Mean<br>abundance<br>across all<br>samples | Standard<br>Deviation | Standard<br>Error |
| --- | --- | --- | --- | --- | --- | --- | --- | --- | --- |
| 1 | Staphylococcus | 0.4118 | 25 | 92.6 | 23 | 85.2 | 0.43002067 | 0.424574076 | 0.0422467 |
| 2 | Pseudomonadaceae_uncl | 0.4077 | 25 | 92.6 | 21 | 77.8 | 0.348833988 | 0.41211472 | 0.041006947 |
| 3 | Bordetella | 0.043 | 15 | 55.6 | 10 | 37 | 0.069704213 | 0.230869672 | 0.022972391 |
| 4 | Burkholderia | 0.0275 | 8 | 29.6 | 4 | 14.8 | 0.038078581 | 0.163958354 | 0.016314466 |
| 5 | Streptococcus | 0.0146 | 19 | 70.4 | 13 | 48.1 | 0.01817556 | 0.107603503 | 0.010706949 |
| 6 | Fusobacterium | 0.01 | 6 | 22.2 | 2 | 7.4 | 0.002678309 | 0.022560873 | 0.002244891 |
| 7 | Stenotrophomonas | 0.0098 | 12 | 44.4 | 6 | 22.2 | 0.020840547 | 0.10566466 | 0.010514027 |
| 8 | Actinobacteria_uncl | 0.0093 | 13 | 48.1 | 8 | 29.6 | 0.020388974 | 0.116331921 | 0.011575459 |
| 9 | Pasteurellaceae_uncl | 0.0085 | 10 | 37 | 6 | 22.2 | 0.002338915 | 0.022733269 | 0.002262045 |
| 10 | Bacteroides | 0.0084 | 11 | 40.7 | 7 | 25.9 | 0.002373978 | 0.022598338 | 0.002248619 |
| 11 | Mycobacterium | 0.0069 | 3 | 11.1 | 2 | 7.4 | 0.006255698 | 0.053828728 | 0.005356159 |
| 12 | Neisseria | 0.0066 | 10 | 37 | 6 | 22.2 | 0.001873116 | 0.017635309 | 0.001754779 |
| 13 | Prevotella_7 | 0.0062 | 6 | 22.2 | 4 | 14.8 | 0.00180565 | 0.016410278 | 0.001632884 |
| 14 | Prevotella | 0.0055 | 9 | 33.3 | 6 | 22.2 | 0.003978868 | 0.022749065 | 0.002263617 |
| 15 | Haemophilus | 0.0035 | 6 | 22.2 | 4 | 14.8 | 0.002634651 | 0.019099189 | 0.00190044 |
| 16 | Enterobacteriaceae_uncl | 0.0035 | 11 | 40.7 | 4 | 14.8 | 0.003576617 | 0.018490395 | 0.001839863 |
| 17 | Corynebacterium_1 | 0.0033 | 15 | 55.6 | 9 | 33.3 | 0.007405359 | 0.053400466 | 0.005313545 |
| 18 | Dolosigranulum | 0.0024 | 4 | 14.8 | 2 | 7.4 | 0.004828915 | 0.048269142 | 0.004802959 |
| 19 | Veillonella | 0.0022 | 9 | 33.3 | 4 | 14.8 | 0.000647501 | 0.005971301 | 0.000594167 |
| 20 | Leptotrichia | 0.0017 | 5 | 18.5 | 2 | 7.4 | 0.000453501 | 0.004464456 | 0.00044423 |
| 21 | Alloprevotella | 0.0008 | 2 | 7.4 | 1 | 3.7 | 0.000202502 | 0.002025486 | 0.000201543 |
| 22 | Gemella | 0.0007 | 4 | 14.8 | 3 | 11.1 | 0.000202893 | 0.001922157 | 0.000191262 |
| 23 | Lachnoanaerobaculum | 0.0005 | 3 | 11.1 | 1 | 3.7 | 0.000147851 | 0.001463586 | 0.000145632 |
| 24 | Peptoniphilus | 0.0005 | 3 | 11.1 | 2 | 7.4 | 0.001928542 | 0.014726115 | 0.001465303 |
| 25 | Selenomonas_3 | 0.0003 | 2 | 7.4 | 1 | 3.7 | 8.75489E-05 | 0.000857603 | 8.53347E-05 |
| 26 | Capnocytophaga | 0.0003 | 7 | 25.9 | 1 | 3.7 | 0.000212675 | 0.001449359 | 0.000144217 |
| 27 | Prevotella_6 | 0.0003 | 4 | 14.8 | 2 | 7.4 | 0.00010341 | 0.000813092 | 8.09056E-05 |
| 28 | Campylobacter | 0.0003 | 4 | 14.8 | 3 | 11.1 | 0.000331682 | 0.001969071 | 0.00019593 |
| 29 | Stomatobaculum | 0.0003 | 3 | 11.1 | 1 | 3.7 | 7.58472E-05 | 0.000724581 | 7.20985E-05 |
| 30 | Anaerococcus | 0.0003 | 7 | 25.9 | 3 | 11.1 | 0.000557374 | 0.003150702 | 0.000313507 |
| 31 | Prevotellaceae_uncl | 0.0002 | 8 | 29.6 | 2 | 7.4 | 0.000517544 | 0.004888722 | 0.000486446 |
| 32 | Pseudomonas | 0.0002 | 12 | 44.4 | 5 | 18.5 | 0.002669558 | 0.02513086 | 0.002500614 |
| 33 | Acinetobacter | 0.0002 | 15 | 55.6 | 7 | 25.9 | 0.0004655 | 0.002066011 | 0.000205576 |
| 34 | Moraxella | 0.0002 | 10 | 37 | 6 | 22.2 | 0.000133847 | 0.000572013 | 5.69174E-05 |
| 35 | Megasphaera | 0.0002 | 4 | 14.8 | 1 | 3.7 | 4.51189E-05 | 0.000428852 | 4.26723E-05 |
| 36 | Granulicatella | 0.0001 | 6 | 22.2 | 1 | 3.7 | 0.000164862 | 0.001226152 | 0.000122007 |
| 37 | Gammaproteobacteria_uncl | 0.0001 | 8 | 29.6 | 4 | 14.8 | 7.13E-05 | 0.000275222 | 2.74E-05 |
| 38 | Bergeyella | 0.0001 | 3 | 11.1 | 1 | 3.7 | 2.43348E-05 | 0.000208357 | 2.07323E-05 |
| 39 | Blautia | 0.0001 | 8 | 29.6 | 3 | 11.1 | 0.000678114 | 0.00655322 | 0.00065207 |
| 40 | Chryseobacterium | 0.0001 | 6 | 22.2 | 4 | 14.8 | 8.801E-05 | 0.000541202 | 5.38517E-05 |
| 41 | Sphingomonas | 0.0001 | 11 | 40.7 | 7 | 25.9 | 5.25892E-05 | 0.00019488 | 1.93912E-05 |
| 42 | Pelomonas | 0.0001 | 10 | 37 | 5 | 18.5 | 0.000100349 | 0.000468823 | 4.66497E-05 |
| 43 | Turicella | 0.0001 | 6 | 22.2 | 2 | 7.4 | 0.000159799 | 0.001021605 | 0.000101654 |
| 44 | Oribacterium | 0.0001 | 2 | 7.4 | 2 | 7.4 | 1.91712E-05 | 0.000151526 | 1.50774E-05 |
| 45 | Neisseriaceae_uncl | 0.0001 | 5 | 18.5 | 2 | 7.4 | 4.27E-05 | 0.000268764 | 2.67E-05 |
| 46 | Ruminococcus_2 | 0.0001 | 4 | 14.8 | 2 | 7.4 | 0.000556224 | 0.00546117 | 0.000543407 |
| 47 | Lactobacillus | 0.0001 | 7 | 25.9 | 4 | 14.8 | 5.14529E-05 | 0.000259294 | 2.58007E-05 |
| 48 | Finegoldia | 0.0001 | 5 | 18.5 | 2 | 7.4 | 0.000151581 | 0.000924509 | 9.19921E-05 |
| 49 | Rothia | 0 | 9 | 33.3 | 5 | 18.5 | 4.07843E-05 | 0.000131184 | 1.30533E-05 |
| 50 | Aquabacterium | 0 | 4 | 14.8 | 3 | 11.1 | 2.00704E-05 | 0.000150486 | 1.49739E-05 |
| 51 | Bradyrhizobium | 0 | 10 | 37 | 6 | 22.2 | 6.77824E-05 | 0.000255806 | 2.54537E-05 |
| 52 | Micrococcus | 0 | 8 | 29.6 | 2 | 7.4 | 5.24264E-05 | 0.000239931 | 2.3874E-05 |
| 53 | [Eubacterium]_nodatum_grp | 0 | 1 | 3.7 | 1 | 3.7 | 1.03E-05 | 0.000103496 | 1.03E-05 |
| 54 | Bacilli_uncl | 0 | 9 | 33.3 | 5 | 18.5 | 4.71E-05 | 0.000145319 | 1.45E-05 |
| 55 | Comamonadaceae_uncl | 0 | 5 | 18.5 | 4 | 14.8 | 7.56E-05 | 0.000504326 | 5.02E-05 |
| 56 | Bacteria_uncl | 0 | 8 | 29.6 | 4 | 14.8 | 4.18E-05 | 0.000146741 | 1.46E-05 |
| 57 | Methylobacterium | 0 | 9 | 33.3 | 6 | 22.2 | 4.05889E-05 | 0.000168225 | 1.6739E-05 |
| 58 | Lachnospiraceae_uncl | 0 | 7 | 25.9 | 2 | 7.4 | 0.000196502 | 0.00188629 | 0.000187693 |
| 59 | Dialister | 0 | 2 | 7.4 | 2 | 7.4 | 8.78323E-05 | 0.000525151 | 5.22544E-05 |
| 60 | Alishewanella | 0 | 4 | 14.8 | 1 | 3.7 | 1.1186E-05 | 8.23999E-05 | 8.1991E-06 |
| 61 | Actinomyces | 0 | 3 | 11.1 | 1 | 3.7 | 1.017E-05 | 6.53118E-05 | 6.49877E-06 |
| 62 | unknown_uncl | 0 | 3 | 11.1 | 1 | 3.7 | 3.06E-05 | 0.000176293 | 1.75E-05 |
| 63 | Subdoligranulum | 0 | 5 | 18.5 | 1 | 3.7 | 0.000167339 | 0.001415227 | 0.00014082 |
| 64 | Porphyromonas | 0 | 5 | 18.5 | 2 | 7.4 | 2.41443E-05 | 0.000159913 | 1.5912E-05 |
| 65 | Hymenobacter | 0 | 5 | 18.5 | 3 | 11.1 | 2.10241E-05 | 0.000105945 | 1.05419E-05 |
| 66 | Alloicoccus | 0 | 6 | 22.2 | 3 | 11.1 | 2.14639E-05 | 0.00012783 | 1.27196E-05 |
| 67 | Domibacillus | 0 | 1 | 3.7 | 1 | 3.7 | 3.03339E-05 | 0.000304852 | 3.03339E-05 |
| 68 | Delftia | 0 | 6 | 22.2 | 1 | 3.7 | 1.54631E-05 | 7.56312E-05 | 7.52559E-06 |
| 69 | Ruminococcaceae_UCG_002 | 0 | 5 | 18.5 | 2 | 7.4 | 8.17E-05 | 0.000513844 | 5.11E-05 |
| 70 | Prevotella_9 | 0 | 7 | 25.9 | 2 | 7.4 | 7.46306E-05 | 0.000568666 | 5.65844E-05 |
| 71 | Prevotella_2 | 0 | 4 | 14.8 | 0 | 0 | 9.40291E-06 | 6.42449E-05 | 6.3926E-06 |
| 72 | Janibacter | 0 | 2 | 7.4 | 2 | 7.4 | 9.20962E-06 | 5.64448E-05 | 5.61646E-06 |
| 73 | Alcaligenaceae_uncl | 0 | 3 | 11.1 | 3 | 11.1 | 2.89E-05 | 0.000109037 | 1.08E-05 |
| 74 | Pseudobutyrvibrio | 0 | 6 | 22.2 | 3 | 11.1 | 0.000103679 | 0.000894836 | 8.90395E-05 |
| 75 | Faecalibacterium | 0 | 6 | 22.2 | 2 | 7.4 | 5.55362E-05 | 0.000342782 | 3.41081E-05 |
| 76 | Sporobacter | 0 | 1 | 3.7 | 1 | 3.7 | 5.71651E-06 | 5.74502E-05 | 5.71651E-06 |
| 77 | Azomonas | 0 | 7 | 25.9 | 2 | 7.4 | 1.67769E-05 | 5.25508E-05 | 5.229E-06 |
| 78 | Anaerostipes | 0 | 3 | 11.1 | 2 | 7.4 | 0.000132057 | 0.001290912 | 0.000128451 |
| 79 | Micrococcaceae_uncl | 0 | 6 | 22.2 | 1 | 3.7 | 1.69E-05 | 8.91E-05 | 8.87E-06 |
| 80 | Fusicatenibacter | 0 | 1 | 3.7 | 1 | 3.7 | 0.000138337 | 0.001390272 | 0.000138337 |
| 81 | Sorangium | 0 | 4 | 14.8 | 1 | 3.7 | 2.22701E-05 | 0.000139415 | 1.38723E-05 |
| 82 | Bacillales_uncl | 0 | 9 | 33.3 | 1 | 3.7 | 1.78E-05 | 8.07E-05 | 8.03E-06 |
| 83 | Xanthomonadaceae_uncl | 0 | 4 | 14.8 | 1 | 3.7 | 7.95E-06 | 4.95E-05 | 4.93E-06 |
| 84 | Staphylococcaceae_uncl | 0 | 7 | 25.9 | 1 | 3.7 | 1.39E-05 | 5.18E-05 | 5.15E-06 |
| 85 | Massilia | 0 | 3 | 11.1 | 1 | 3.7 | 1.43407E-05 | 7.2316E-05 | 7.19571E-06 |
| 86 | Abiotrophia | 0 | 5 | 18.5 | 1 | 3.7 | 9.92484E-06 | 3.74956E-05 | 3.73095E-06 |
| 87 | Cloacibacterium | 0 | 4 | 14.8 | 3 | 11.1 | 1.59258E-05 | 9.49309E-05 | 9.44598E-06 |
| 88 | Ruminococcus_1 | 0 | 1 | 3.7 | 1 | 3.7 | 9.22517E-05 | 0.000894035 | 8.89598E-05 |
| 89 | Actinobacillus | 0 | 1 | 3.7 | 0 | 0 | 2.94234E-06 | 2.95702E-05 | 2.94234E-06 |
| 90 | Facklamia | 0 | 2 | 7.4 | 1 | 3.7 | 5.64365E-06 | 4.08389E-05 | 4.06363E-06 |
| 91 | Devosia | 0 | 2 | 7.4 | 1 | 3.7 | 5.64365E-06 | 4.08389E-05 | 4.06363E-06 |
| 92 | Brevundimonas | 0 | 3 | 11.1 | 1 | 3.7 | 1.62809E-05 | 0.00011187 | 1.11315E-05 |
| 93 | Ruminococcaceae_uncl | 0 | 2 | 7.4 | 1 | 3.7 | 2.00E-05 | 0.000140293 | 1.40E-05 |
| 94 | Alphaproteobacteria_uncl | 0 | 6 | 22.2 | 2 | 7.4 | 1.72E-05 | 9.01E-05 | 8.97E-06 |
| 95 | Tepidiphilus | 0 | 2 | 7.4 | 1 | 3.7 | 1.95942E-05 | 0.000186642 | 1.85716E-05 |
| 96 | Conchiformibius | 0 | 1 | 3.7 | 1 | 3.7 | 4.57391E-06 | 4.59672E-05 | 4.57391E-06 |
| 97 | Candidate_division_SR1_uncl | 0 | 2 | 7.4 | 0 | 0 | 3.12E-06 | 2.21E-05 | 2.20E-06 |
| 98 | Sphingomonadales_uncl | 0 | 6 | 22.2 | 0 | 0 | 3.76E-05 | 0.000236781 | 2.36E-05 |
| 99 | Lachnoclostridium | 0 | 8 | 29.6 | 3 | 11.1 | 3.29414E-05 | 0.00020429 | 2.03276E-05 |
| 100 | Rheinheimera | 0 | 3 | 11.1 | 0 | 0 | 7.04733E-06 | 4.60856E-05 | 4.58569E-06 |

**Supplemental table 3.**
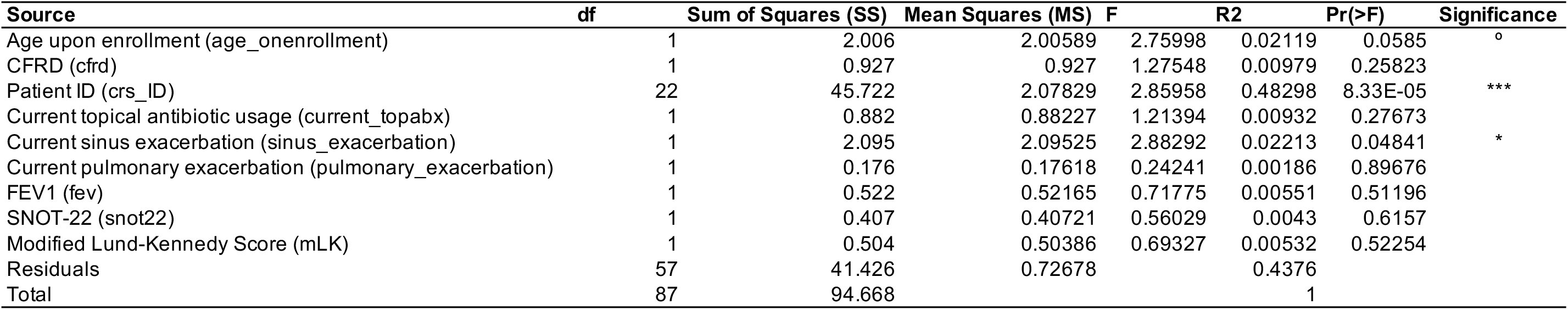
CF CRS microbial communities are highly individualized, but may share similarities during sinus exacerbation. PERMANOVA results describing the proportion of variance in sample composition attributable to variables tested (“source” of variation). Two sources contributed a significant amount of variance (p <0.05; Patient ID and whether or not a study participant was experiencing a sinus exacerbation), whereas enrollment age has a non-statistically significant effect (p < 0.1) and the remaining variables had non-significant effects (p > 0.1). The name of the variable as it appears in the metadata sheet is included in parentheses in the first column (“Source”). Terms were added sequentially and the model was run with 12000 permutations. Significance levels were determined by the Pr(>F). ***: p <0.001, **: p < 0.01, *: p < 0.05, ° : p < 0.1, blank: p >0.1.

**Supplemental table 4.**
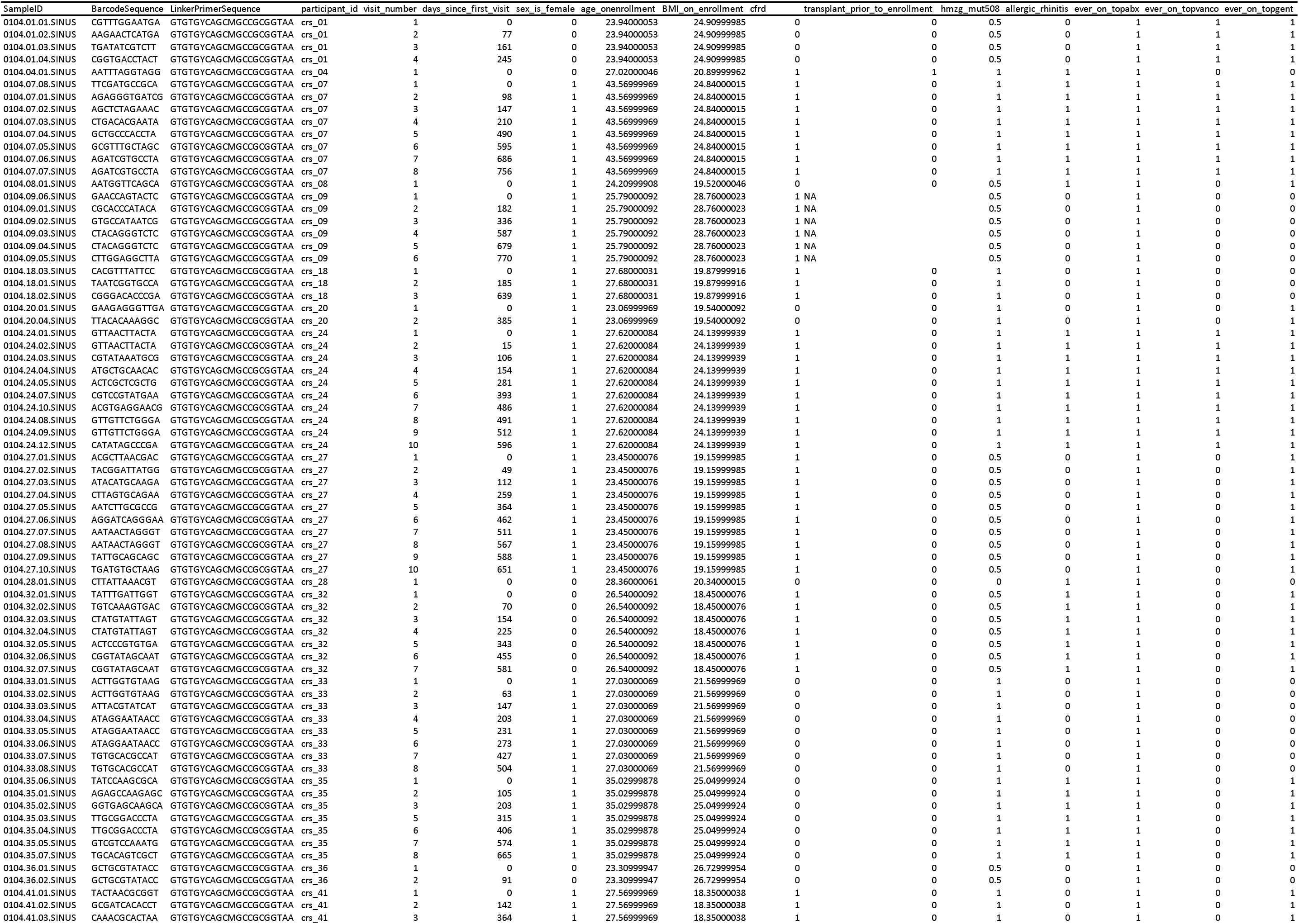
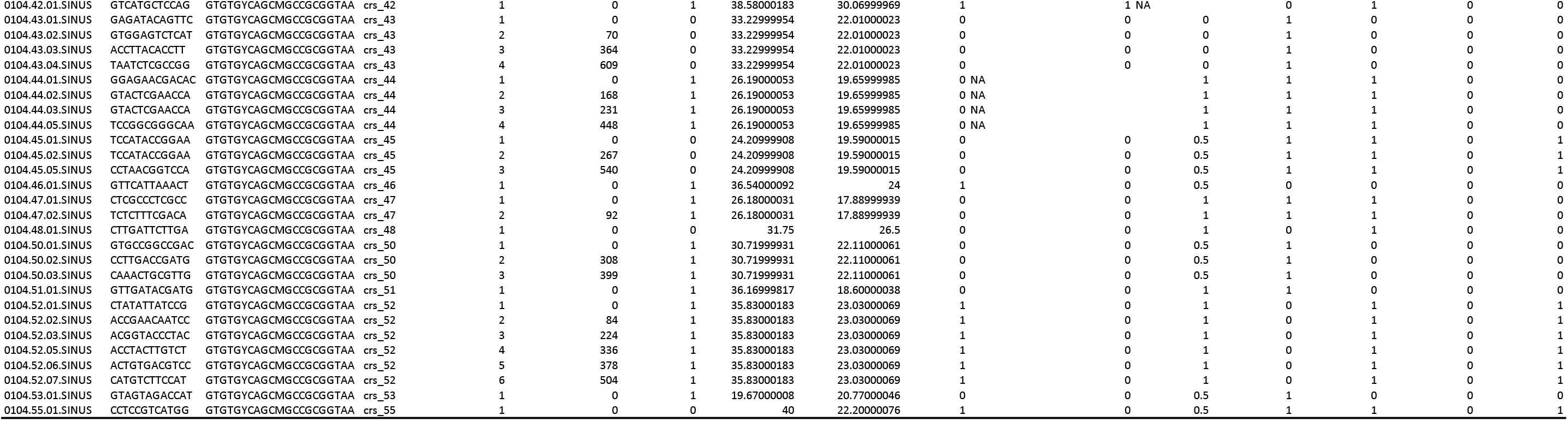
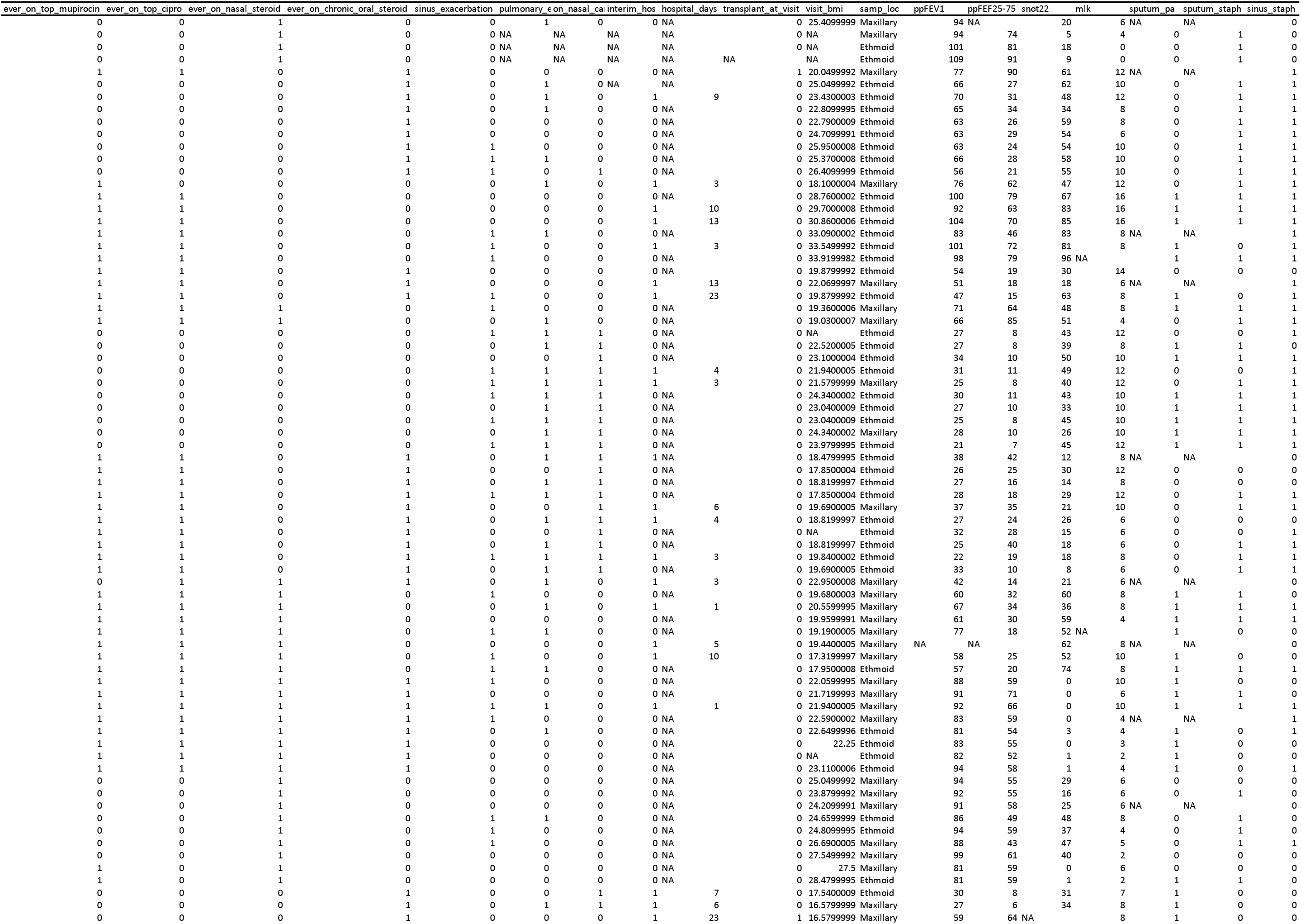
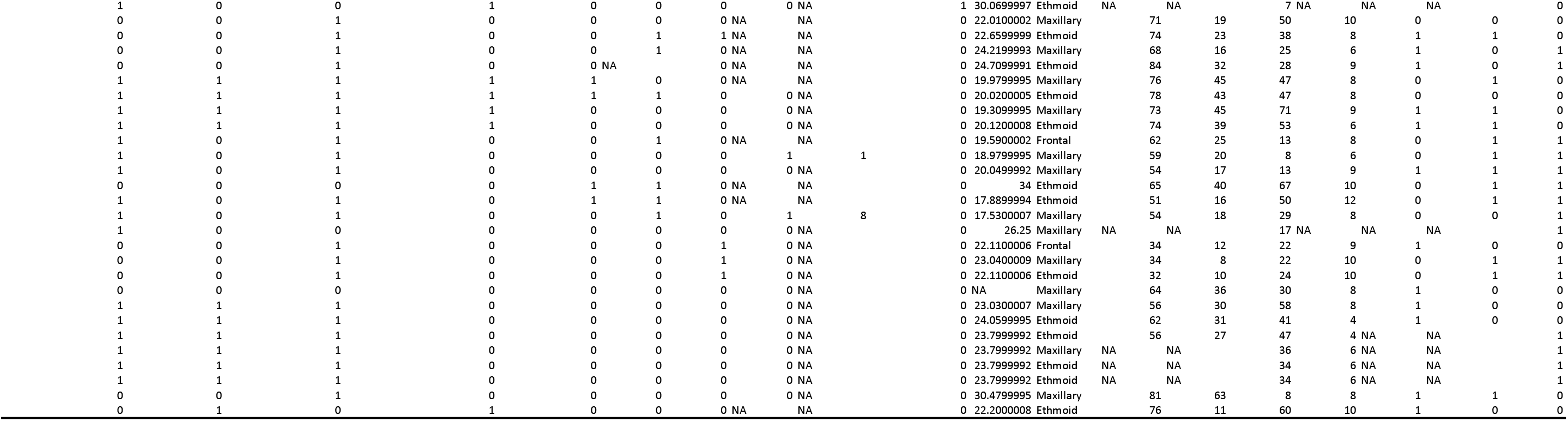
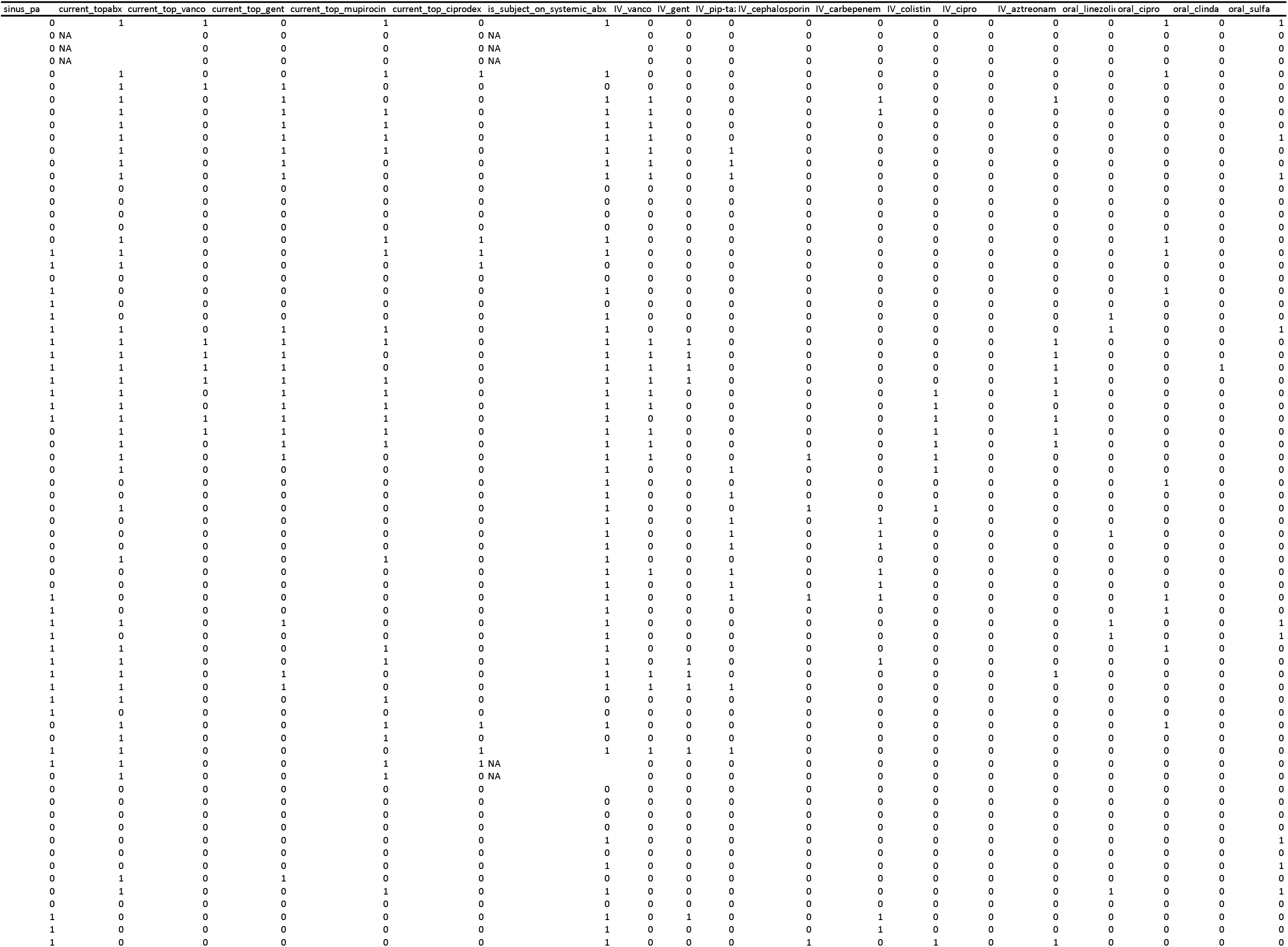
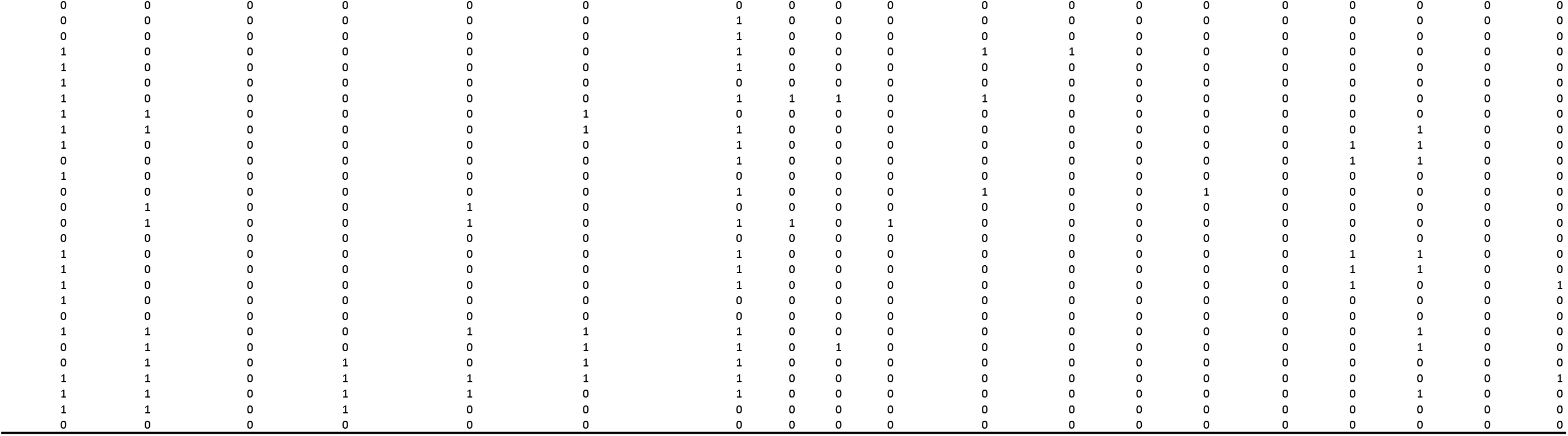
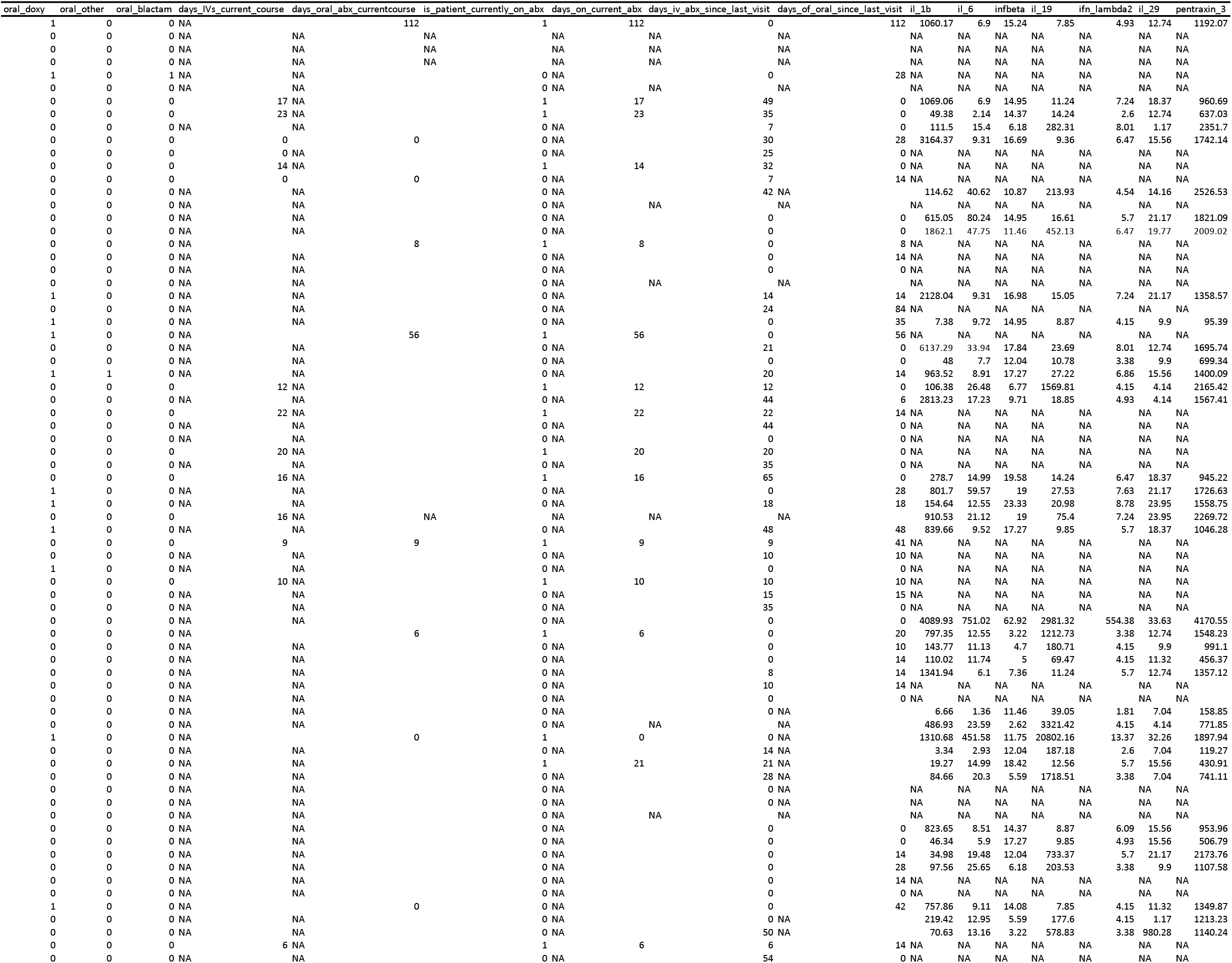
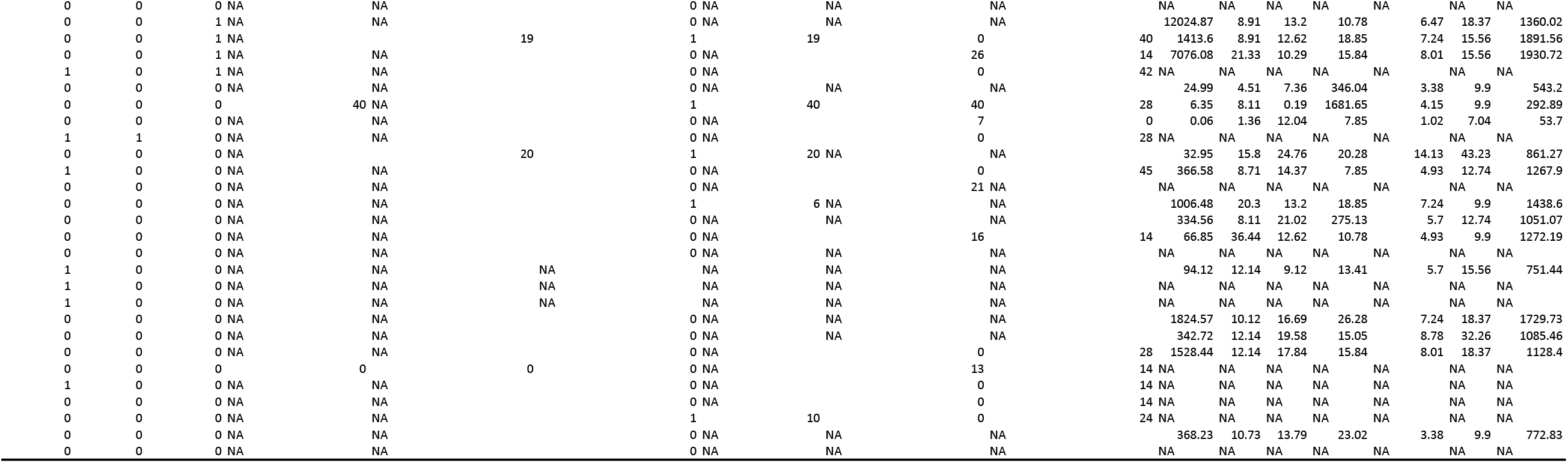
Metadata associated with each sequenced microbiota sample. See the codebook in Supplemental table 5 for a description of each variable.

**Supplemental table 5.**
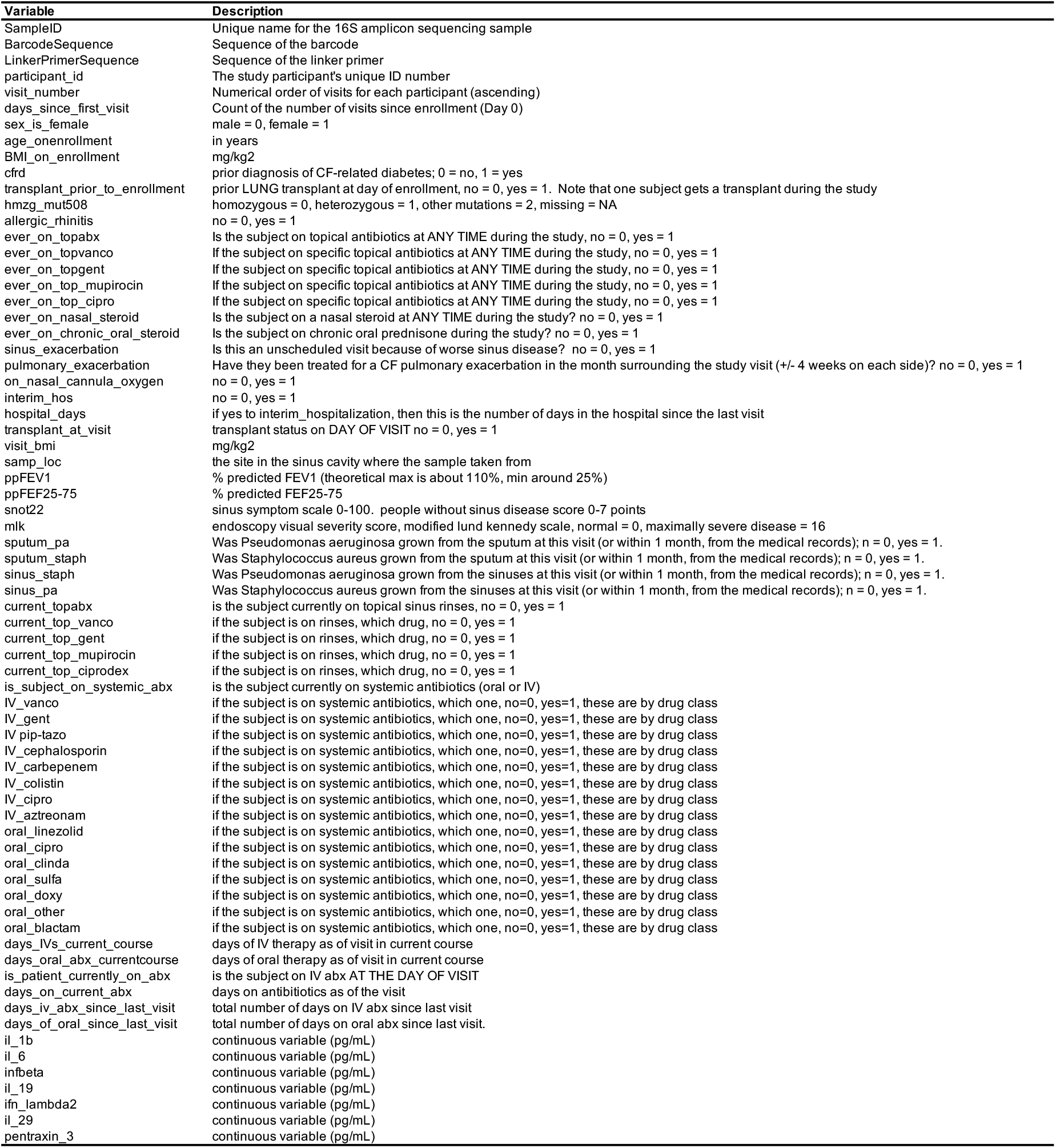
Codebook describing each variable in the metadata. See Supplemental table 4 for the metadata.

